# MEF2C phosphorylation is required for chemotherapy resistance in acute myeloid leukemia

**DOI:** 10.1101/107201

**Authors:** Fiona C. Brown, Eric Still, Paolo Cifani, Sumiko Takao, Casie Reed, Scott B. Ficarro, Richard P. Koche, Peter Romanienko, Willie Mark, Conor O’Donnell, Barbara Spitzer, Crystal Stutzke, Andrei V. Krivtsov, Gayle Pouliot, Nathanael Gray, Jarrod A. Marto, Scott Armstrong, Alex Kentsis

## Abstract

**HIGHLIGHTS:**

- MEF2C S222 phosphorylation is a specific marker of chemotherapy resistance in diagnostic AML patient specimens.
- MEF2C S222 phosphorylation is dispensable for normal hematopoiesis in mice, as established using genome editing *in vivo*, but is required for MLL-AF9 induced leukemogenesis.
- MARK kinases specifically phosphorylate MEF2C S222, potentiating its transcriptional activity.
- Chemical inhibition of MARK-induced MEF2C phosphorylation overcomes chemotherapy resistance of and exhibits selectivity toxicity against MEF2C-activated human AML cells.

**SUMMARY:** In acute myeloid leukemia, chemotherapy resistance remains prevalent and poorly understood. Using functional proteomics of patient AML specimens, we identified MEF2C S222 phosphorylation as a specific marker of primary chemoresistance. We found that *Mef2c*^*S222A/S222A*^ knock-in mutant mice engineered to block MEF2C phosphorylation exhibited normal hematopoiesis, but were resistant to leukemogenesis induced by *MLL-AF9*. MEF2C phosphorylation was required for leukemia stem cell maintenance, and induced by MARK kinases in cells. Treatment with the selective MARK inhibitor MRT199665 caused apoptosis of MEF2C-activated human AML cell lines and primary patient specimens, but not those lacking MEF2C phosphorylation. These findings identify kinase-dependent dysregulation of transcription factor control as a determinant of therapy response in AML, with immediate potential for improved diagnosis and therapy for this disease.

## INTRODUCTION

Acute myeloid leukemias (AML) are cancers of the blood that originate in the hematopoietic progenitor cells as a result of the accumulation of genetic mutations that lead to cell transformation. Recent advances in genomic profiling have revealed distinct genetic subsets of AML, including specific mutational classes of cytogenetically normal and chromosomally-rearranged leukemias (Cancer Genome Atlas Research, 2013; Papaemmanuil et al., 2016; Schuback et al., 2013). Overall, AML is characterized by a prevalence of mutations of genes encoding regulators of gene expression, such as *MLL-AF9* (*KMT2A-MLLT3*) gene fusion that dysregulates the expression of genes controlling self-renewal, differentiation and cell survival (Zuber et al., 2011). Recent studies have begun to reveal specific molecular dependencies that can be used for improved targeted therapies in AML (Coombs et al., 2016). In spite of this knowledge, intensive chemotherapy and stem cell transplantation continue to be essential means to achieve cure in the treatment of AML. However, current chemotherapy regimens remain inadequate and fail to induce or sustain remissions in more than 50% of adults and 30% of children with AML (Breems et al., 2005; Burnett et al., 2011; de Rooij et al., 2015). Thus, improved therapeutic strategies to overcome or block chemotherapy resistance are needed.

Patients with distinct molecularly identifiable types of leukemias have been found to exhibit varying degrees of response to chemotherapy, leading to therapy intensification for patients with high-risk disease (Ding et al., 2012; Farrar et al., 2016; Kihara et al., 2014; Klco et al., 2015). However, human leukemias also exhibit large heterogeneity of chemotherapy response, and its molecular determinants continue to remain poorly understood. For example, mutations of *DNMT3A*, as well as gene fusions such as *MLL-AF9* and *NUP98-NSD1,* have been found to contribute to chemotherapy resistance (Guryanova et al., 2016; Hollink et al., 2011; Zuber et al., 2009). However, patients with leukemias with these and other mutations can nonetheless achieve durable remissions (Papaemmanuil et al., 2016). Likewise, leukemias that exhibit primary chemotherapy resistance that is refractory to induction chemotherapy do not show significant enrichment for these or other single gene mutations thus far (Brown et al., 2017; Farrar et al., 2016; Klco et al., 2015). Thus, additional genetic or molecular mechanisms must cause chemotherapy resistance in AML.

In support of this notion, human leukemias and their genetically-engineered mouse models are characterized by distinct cell populations, comprising defined functional compartments such as leukemia stem or initiating cells that exhibit unique phenotypic properties including self-renewal and enhanced cell survival (Hope et al., 2004; Somervaille and Cleary, 2006). In part, these behaviors are caused by co-option of developmental programs that regulate normal hematopoiesis, such as for example self-renewal and therapy resistance of *MLL-AF9* leukemias (Krivtsov et al., 2006; Zuber et al., 2011). In particular, MEF2C, a member of the MADS family of transcription factors, normally regulates hematopoietic self-renewal and differentiation, supports the proliferation of *MLL-AF9* leukemias, and is associated with increased risk of relapse when highly expressed in multiple subtypes of AML in patients (Herglotz et al., 2016; Krivtsov et al., 2006; Laszlo et al., 2015; Schuler et al., 2008; Schwieger et al., 2009; Stehling-Sun et al., 2009; Wang et al., 2016). The precise molecular mechanisms by which MEF2C is dysregulated in AML are not known. However, recurrent mutations of other MEF2 family members, including *MEF2B*, *MEF2C* and *MEF2D* in refractory lymphoblastic leukemias and lymphomas (Gu et al., 2016; Pon et al., 2015; Ying et al., 2013), suggest that MEF2C may regulate a general mechanism of therapy resistance.

Here, using recently developed high-accuracy mass spectrometry techniques, we determined phospho-signaling profiles of human AML specimens collected at diagnosis from patients with primary chemotherapy resistance and failure of induction chemotherapy. Analysis of these profiles revealed high levels of phosphorylation of S222 of MEF2C, which was found to be significantly associated with primary chemotherapy resistance in an independent cohort of cytogenetically normal and MLL-rearranged leukemias. By integrating genome editing, biochemical and cell biological approaches, we tested the hypothesis that MEF2C phosphorylation promotes chemotherapy resistance and that its blockade can be leveraged for improved AML therapy. These studies have revealed an unexpected dependence on kinase-dependent dysregulation of transcription factor control as a determinant of therapy response in AML, with immediate potential for translation into improved diagnosis and therapy for this disease.

## Results

### Phosphorylation of S222 in MEF2C is a specific marker of AML chemotherapy resistance

Previously, we assembled a cohort of primary AML specimens collected at diagnosis from patients with failure of induction chemotherapy and those who achieved remission, after two cycles of cytarabine and daunorubicin-based induction chemotherapy, matched for AML subtypes and therapy (Brown et al., 2017). In this analysis, we found that defined gene mutations were associated with primary chemotherapy resistance only in a minority of cases. Thus, we sought to investigate alternative molecular mechanisms that may explain primary chemotherapy resistance in AML.

We focused on phosphosignaling because activation of kinases is one of the hallmarks of AML pathogenesis (Kentsis et al., 2012; Zheng et al., 2004). Recent advances in quantitative proteomics, particularly in high-efficiency, multi-dimensional fractionation platforms (Ficarro et al., 2011) enable in-depth analysis of signaling molecules from rare cell populations (Lu et al., 2014). Leukemia cells purified from a discovery cohort of eight diagnostic AML bone marrow aspirate specimens were analyzed by metal affinity chromatography (IMAC) (Ficarro et al., 2009) and isobaric tagging (iTRAQ) mass spectrometry (Choe et al., 2007). This yielded 2,553 unique phosphopeptides, 53 of which were significantly enriched in induction failure specimens (Data S1, Figure S1A). We identified phosphorylation of serine 222 (pS222) in MEF2C as being the most highly abundant phosphoprotein in induction failure specimens as compared to age, therapy, and disease-matched remission specimens (*p* = 5.0 × 10^-3^, t-test, Figures 1A and S1B). The observed MEF2C pS222 peptide was distinct from the related MEF2A, MEF2B, and MEF2D proteins (Figure S1C). In addition, this analysis revealed phosphoproteins previously implicated in therapy resistance, such as HSBP1 pS15 (Kasimir-Bauer et al., 1998), as well as those not previously observed but likely functional such as HGF pT503 (Kentsis et al., 2012).

To investigate the diagnostic significance of MEF2C pS222 in primary AML chemotherapy resistance, we assessed the prevalence of MEF2C pS222 in an independent cohort of 29 pediatric and adult primary AML specimens matched for age, disease biology and therapy (Table S1) (Brown et al., 2017). As previously observed, this cohort included specimens with gene mutations associated with poor prognosis such as those with cryptic rearrangements of *MLL/KMT2A* and combined *DNMT3A* and *NPM1* mutations (Table S1). We developed an affinity-purified antibody against MEF2C pS222, and validated its specificity using synthetic peptide competition and phosphatase assays (Figure S2A-C). We found that expression of MEF2C pS222 and total MEF2C were significantly associated with induction failure and primary chemotherapy resistance (*p* = 5.2 × 10^−3^ and 1.6 − 10^−3^, respectively; Figures 1B-C and S2D-E). *Post hoc* review of four specimens with moderate MEF2C pS222 expression from remission cases at diagnosis revealed that these cases had in fact early relapse after initial induction therapy treatment (Figure 1C), corroborating the specific association of MEF2C pS222 with AML chemotherapy resistance. Additionally, three of these specimens had *MLL/KMT2A* gene rearrangements (Figure 1C). Thus, MEF2C pS222 is a specific marker of primary chemoresistance and failure of induction therapy in AML.

**Figure 1.**
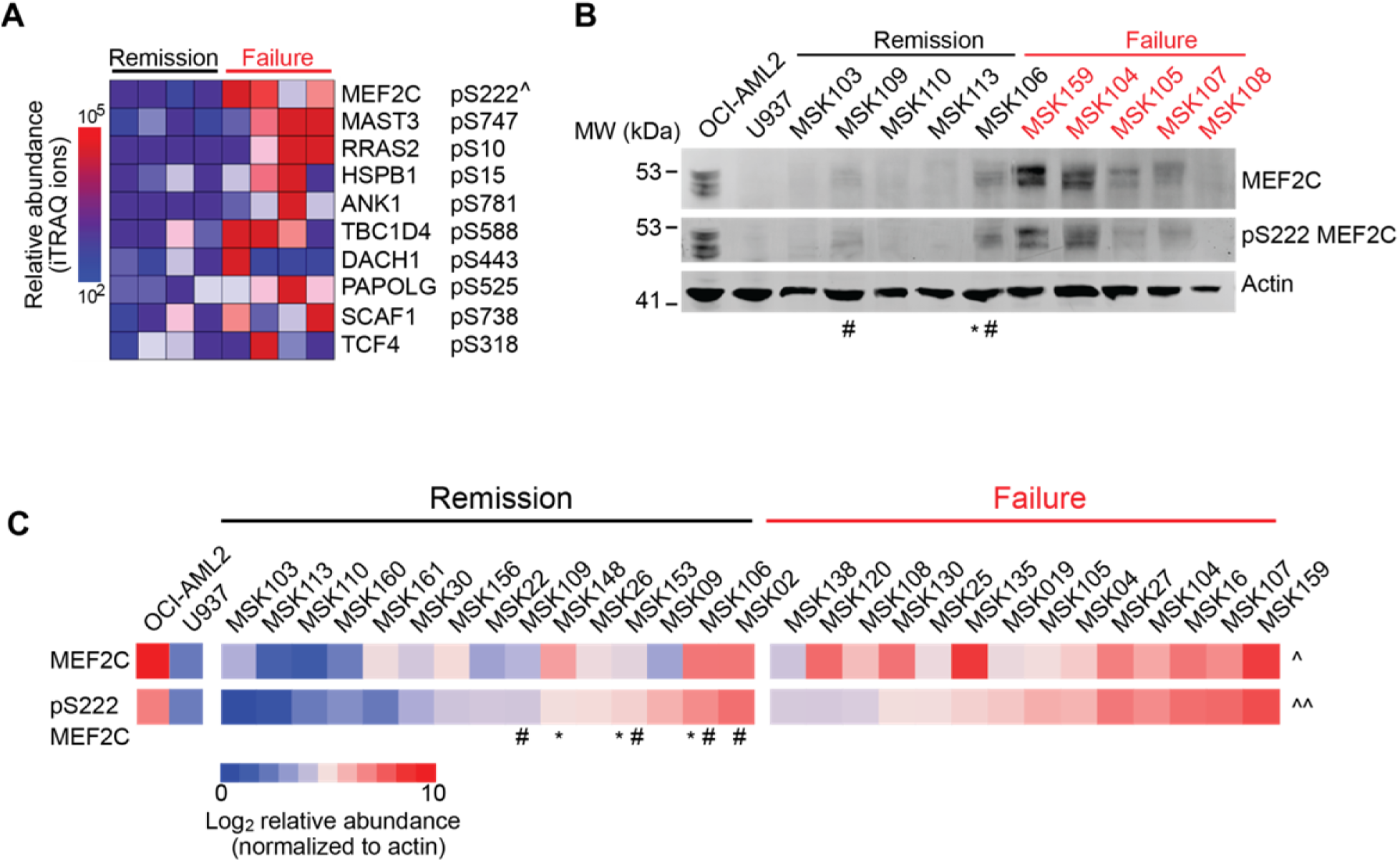
Phosphorylation of MEF2C at serine 222 is associated with primary AML chemoresistance. (A) Heatmap of the 10 top differentially abundant protein phosphorylation sites detected in diagnostic AML specimens in 4 patients with primary chemotherapy resistance and induction failure, as compared to 4 age, disease and therapy-matched patients who achieved complete induction remission. Relative abundance is represented by the blue-to-red color gradient of iTRAQ reporter ion intensities. ^ *p* = 5.0 × 10^−3^ (B) Representative Western immunoblot analysis for MEF2C and pS222 MEF2C in a cohort of age, disease and therapy-matched AML patient specimens with induction failure and complete remission. The human AML cell lines OCI-AML2 and U937 serve as positive and negative controls for MEF2C expression and S222 phosphorylation, respectively. (C) Heatmap of MEF2C expression and S222 phosphorylation in a matched cohort of 29 specimens, as measured using quantitative fluorescent immunoblotting, and normalized to actin. ^ and ^^ *p* = 1.6 × 10^−3^ and 5.2 × 10^−3^ for remission versus failure for MEF2C and pS222 MEF2C respectively (t-test). ^#^ and * denote specimens from patients who achieved complete remission but experienced AML relapse, and those with cryptic rearrangement of *MLL/KMT2A*, respectively, as additionally explained in Figure S3.

### MEF2C pS222 is dispensable for normal hematopoiesis

The activity of MEF2C is known to be regulated by post-translational modifications, including acetylation, sumoylation, and phosphorylation, that can affect the recruitment of transcriptional co-repressors and co-activators (Kang et al., 2006; Ma et al., 2005; McKinsey et al., 2002). Analysis of existing phosphosignaling data revealed the presence of MEF2C S222 phosphorylation in human K562 leukemia cells and cytokine-stimulated hematopoietic progenitor cells (Ficarro et al., 2014; Zhou et al., 2013). The specific association of MEF2C pS222 with failure of induction chemotherapy raises the possibility that MEF2C S222 phosphorylation promotes chemotherapy resistance in AML. To test this hypothesis, we first sought to investigate the potential function of MEF2C S222 phosphorylation in normal hematopoiesis. Thus, we engineered knock-in mice harboring a loss-of-function *Mef2c S222A* allele that cannot be phosphorylated, and a gain-of-function *S222D* allele that mimics phospho-serine by using CRISPR/Cas9 genome editing (Figure 2A, S3A). Genotyping of founder animals from both *Mef2c*^*S222A*^ and *Mef2c*^*S222D*^ strains identified the mutant alleles in 10% of born mice. We confirmed the absence of off-target mutations of the *Mef2c* locus by genomic DNA sequencing of each founder animal. To control for possible off-target effects, we obtained two independent founder strains for both *Mef2c*^*S222A*^ and *Mef2c*^*S222D*^ alleles, and backcrossed them to wild-type C57BL/6J mice. Subsequently, *Mef2c*^*S222A/S222A*^ and *Mef2c*^*S222D/S222D*^ mice obtained from heterozygous intercrosses were detected at expected Mendelian ratios (Figure S3B), and exhibited approximately equal Mef2c protein expression, as measured by Western immunoblotting of B220+ bone marrow cells, which are known to exhibit high Mef2c expression (Figure 3B). *Mef2c*^*S222A/S222A*^ and *Mef2c*^*S222D/S222D*^ animals exhibited normal growth as compared to wild-type litter mates (Figure S3C and S3D), and had no discernible defects on histologic analysis of tissues where physiologic activities of Mef2c are required for normal development (Li et al., 2008; Lin et al., 1997), including brain, heart, skeletal muscle, and spleen (data not shown).

**Figure 2.**
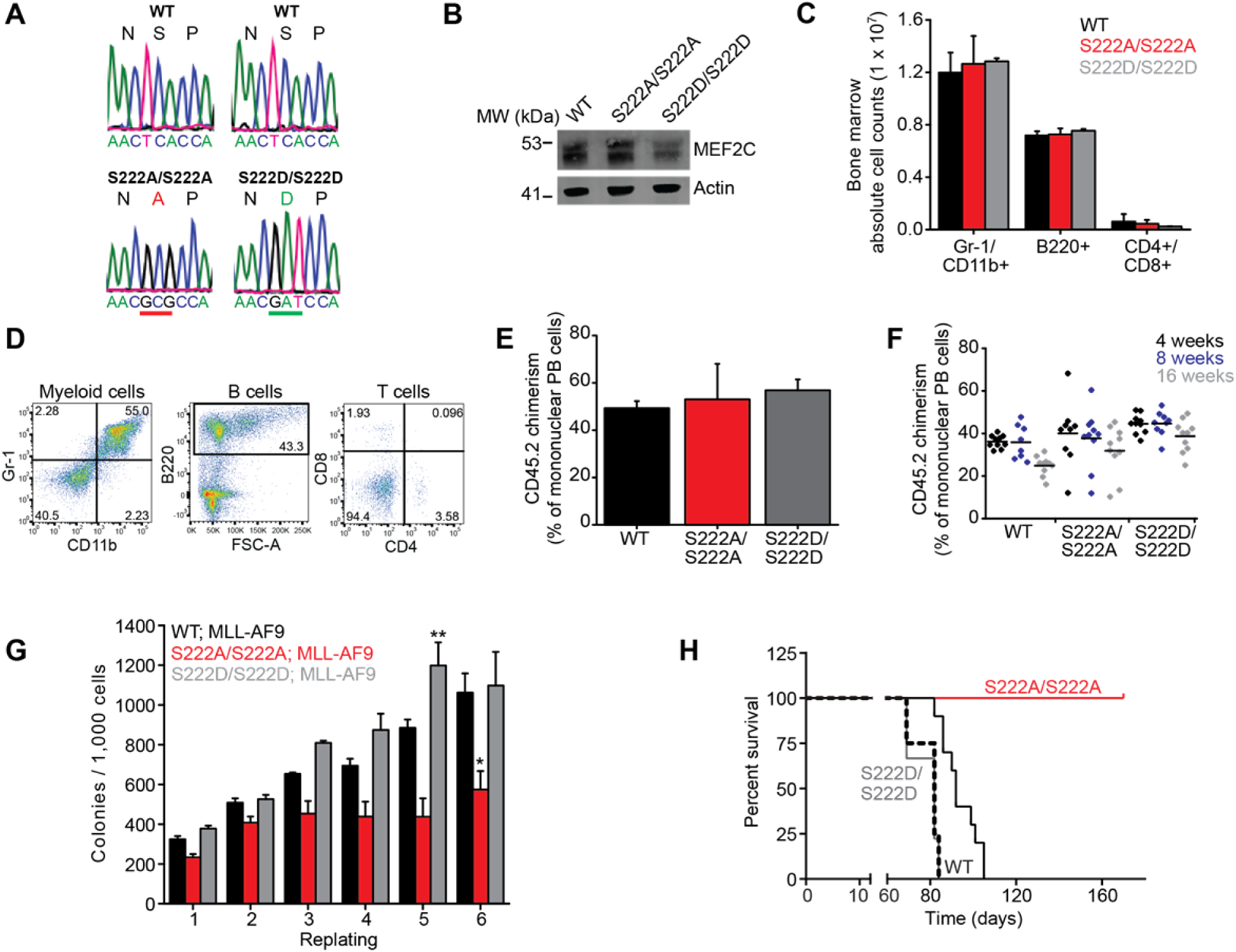
A therapeutic window for targeting MEF2C phosphorylation in AML. (A) Sequencing electropherograms of tail genomic DNA from *Mef2c*^*S222A/S222A*^ and *Mef2c*^*S222D/S222D*^ mice, demonstrating specific CRISPR/Cas9-induced c.TCA>GCG and c.TCA>GAT mutations, as underlined red and green, respectively. (B) Western immunoblot of bone marrow B220+ cells from *Mef2c*^*S222A/S222A*^ and *Mef2c*^*S222D/S222D*^ mice. (C) Total numbers of myeloid (Gr-1^+^/CD11b^+^), B-cell (B220^+^) and T-cell (CD4^+^/CD8^+^) bone marrow cells from *Mef2c*^*S222A/S222A*^ and *Mef2c*^*S222D/S222D*^ mice as assessed by (D) FACS analysis. Error bars represent standard deviation of the mean from 3 mice. (E) Peripheral blood chimerism of CD45.2+ cells at 4 weeks following competitive transplantation. Error bars represent standard deviation of the mean of 10 animals per group. (F) Peripheral blood engraftment of CD45.2+ cells as a function of time post-transplant. Bars denote mean. (G) Serial replating clonogenic efficiencies of bone marrow GMP cells from *Mef2c* ^*S222A/S222A*^, *Mef2c*^*S222D/S222D*^ and wild-type litter mates transduced with *MLL-AF9*. Error bars represent standard deviation of the mean of 3 biological replicates (additional data in). * and ** *p* = 3.3 × 10^−3^ and 1.1 × 10^−2^ of *WT; MLL-AF9* versus *S222A/S222A; MLL-AF9* and *S222D/S222D; MLL-AF9,* respectively (t-test). (H) Kaplan–Meier survival curves of mice transplanted with *MLL-AF9* transformed bone marrow GMP cells from *Mef2c*^*S222A/S222A*^, *Mef2c*^*S222D/S222D*^ and wild-type litter mate controls. *p* = 6.8 × 10^−9^ for *Mef2c*^*S222A/S222A*^ vs wild-type litter mates, log rank test for 10 animals per group. Solid and dashed black lines denote wild-type liter mates for *Mef2cS222A/S222A* and *2cS222D/S222D*, respectively.

Consistent with the dispensability of MEF2C S222 phosphorylation for normal hematopoiesis, *Mef2c*^*S222A/S222A*^ and *Mef2c*^*S222D/S222D*^ mice exhibited normal peripheral blood counts and cell morphologies (Figures S4A and S4B). Likewise, we observed no significant differences in the absolute number of mature blood cells from myeloid and lymphoid lineages in the bone marrow of *Mef2c*^*S222A/S222A*^ and *Mef2c*^*S222D/S222D*^ mice, as compared to wild-type litter mates (Figures 2C, 2D and S4C). Similarly, *Mef2c*^*S222A/S222A*^ and *Mef2c*^*S222D/S222D*^ bone marrow granulocyte-macrophage progenitor (GMP) cells exhibited preserved clonogenic capacity as compared to wild-type *Mef2c* GMP cells from litter mate control mice (Figure S4D). Likewise, CD45.2^+^ *Mef2c*^*S222A/S222A*^ or *Mef2c*^*S222D/S222D*^ hematopoietic stem cells displayed preserved repopulation capacity in competitive bone marrow transplants with wild-type CD45.1^+^ cells in lethally irradiated mice, as monitored by CD45.2^+^ / CD45.1^+^ chimerism of peripheral blood cells in recipient mice over 16 weeks post-transplantation as compared to wild-type controls (Figures 2E-F and S4E-F). Thus, MEF2C S222 phosphorylation is dispensable for normal steady-state and stress hematopoiesis.

### MEF2C phosphorylation is required for MLL-AF9 leukemogenesis

Overexpression of *Mef2c* itself is not sufficient to cause leukemia, but potently cooperates with other leukemia oncogenes (Du et al., 2005). In particular, *Mef2c* is required for the maintenance of MLL-rearranged mouse leukemias (Krivtsov et al., 2006; Schwieger et al., 2009). Since we observed high levels of MEF2C pS222 in primary AML specimens with leukemias with cryptic *MLL* rearrangements, including leukemias with *MLL-AF9* fusions, we reasoned that MEF2C phosphorylation may be required for *MLL-AF9*-induced leukemogenesis.

To test this hypothesis, we assessed leukemia induction by retroviral *MLL-AF9* in bone marrow GMP cells of *Mef2c*^*S222A/S222A*^*, Mef2c*^*S222D/S222D*^ or wild-type litter mates. Bone marrow GMP cells were transduced with the *MSCV-IRES-GFP* retrovirus encoding *MLL-AF9*, and GFP-expressing cells were purified by fluorescence-activated cell sorting (FACS) and plated in methylcellulose for clonogenic assays (Figure 2G). We observed a significant reduction of clonogenic efficiency in serial replating of the *Mef2c*^*S222A/S222A*^ progenitor cells transformed with *MLL-AF9*, as compared to wild-type controls (*p* = 3.3 × 10^−3^, t-test, Figure 2G). In contrast, *MLL-AF9*-transformed *Mef2c*^*S222D/S222D*^ progenitor cells exhibited enhanced replating efficiencies (*p* = 1.1 × 10^−2^, t-test, Figure 2G). We observed the same phenotype in clonogenic assays of bone marrow GMP cells from two independent *Mef2c*^*S222A/S222A*^ founder strains (Figure S4G), consistent with the specific effects of *Mef2c S222A*.

To test the effects of MEF2C S222 phosphorylation on leukemia initiation *in vivo*, *MLL-AF9* transduced cells were transplanted into lethally irradiated C57BL/6J mice by intravenous tail vein injection (Figure S4H). Recipient mice transplanted with *MLL-AF9*-transduced cells derived from wild-type litter mate control and *Mef2c*^*S222D/S222D*^ mice became moribund at approximately 80 days post-transplant with more than 90% of the bone marrow replaced with GFP+ leukemia cells (Figures 2H, S4I). In contrast, mice transplanted with *MLL-AF9*-transduced *Mef2c*^*S222A/S222A*^ cells survived for more than 170 days post-transplant without physical signs of leukemia development (*p* = 6.8 × 10^-9^, log-rank test, Figure 2H). We observed no GFP-expressing cells in the peripheral blood and bone marrow of these mice (Figure S4I), consistent with the essential function for MEF2C pS222 in leukemia stem cell homing or engraftment, as supported by prior studies (Schwieger et al., 2009). Thus, MEF2C S222 phosphorylation is required for leukemia initiation by the MLL-AF9 oncogene.

### MEF2C phosphorylation cooperates with MLL-AF9 induced leukemogenesis and is required for leukemia stem cell maintenance

Protection of *Mef2c*^*S222A/S222A*^ hematopoietic progenitor cells from *MLL-AF9*-induced leukemogenesis suggests that MEF2C S222 phosphorylation may be necessary for the survival of MLL-AF9-transformed leukemia cells. To investigate the function of MEF2C S222 phosphorylation in *MLL-AF9*-induced leukemia maintenance, we transformed wild-type bone marrow GMP cells by retroviral transduction of *MSCV-IRES-GFP MLL-AF9*, and co-expressed wild-type *MEF2C* or S222A mutant using the *MSCV-IRES-tdTomato (MIT)* retrovirus (Figure S5A). This allowed us to purify GFP/tdTomato double-positive *MLL-AF9*-transformed cells using FACS where MEF2C transgenes function as dominant mutants because of their dimerization with endogenous *Mef2c* (Mao et al., 1999; Molkentin et al., 1996). We confirmed that wild-type and dominant negative MEF2C S222A were expressed at approximately equal levels by Western immunoblotting, with MEF2C S222A transduced cells exhibiting reduced levels of pS222 as compared to wild-type control (Figure S5B). We found that *MLL-AF9* leukemia cells expressing MEF2C S222A exhibited transient reduction in clonogenic efficiency *in vitro* as compared to cells expressing wild-type MEF2C (Figure S5C), and displayed increased apoptosis (Figure S5D). Notably, when allowed to develop leukemia (Figure S5E), and secondarily transplanted into sub-lethally irradiated mice to assess leukemia maintenance *in vivo*, *MLL-AF9;MEF2C S222A* leukemias were significantly impaired in leukemia development as compared to *MLL-AF9;MEF2C* wild-type leukemias (*p* = 2.2 × 10^-10^, log-rank test, Figure 3A). Using limiting dilution analysis, we determined that *MLL-AF9;MEF2C S222A* leukemias exhibited more than 8-fold reduction in leukemia stem frequency as compared to *MLL-AF9; MEF2C* wild-type leukemias (1/139 versus 1/16, respectively, *p* = 6.7 × 10^-8^, chi-squared test, Figure 3B and S5F-H). These results indicate that MEF2C S222 phosphorylation is required for the maintenance of MLL-AF9 leukemia stem cells *in vivo*.

**Figure 3.**
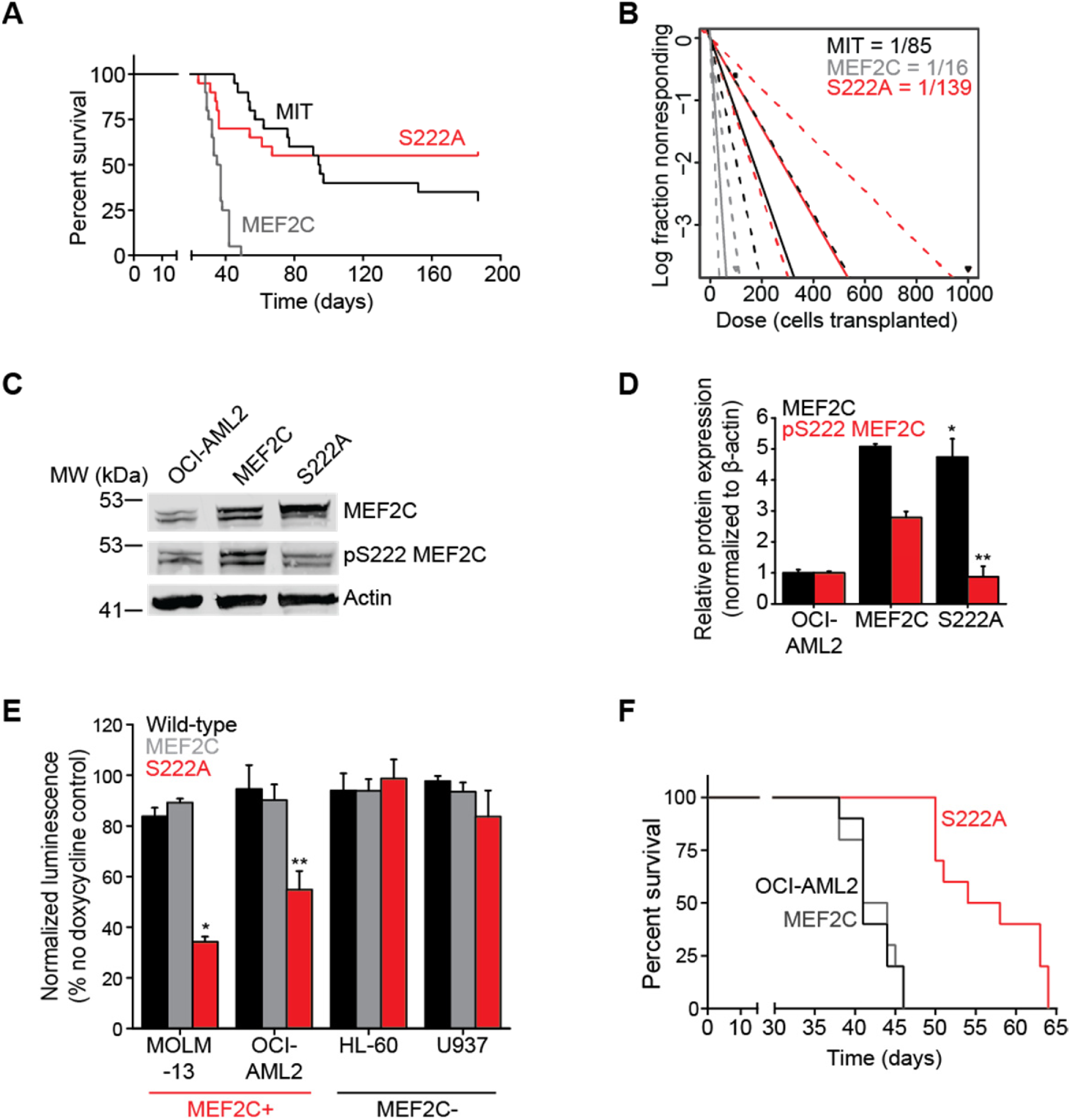
MEF2C phosphorylation is required for leukemia stem cell survival and maintenance in MLL-rearranged leukemia. (A)Kaplan–Meier survival curves of secondary recipient mice transplanted with 100 cells of wild-type *MLL-AF9; MEF2C,* dominant negative *MLL-AF9; MEF2C S222A* orcontrol *MLL-AF9; MIT* transformed leukemias. *p* = 2.2 × 10^−10^ for *MLL-AF9; MEF2C S222A* versus *MLL-AF9; MEF2C*, log rank test for 10 animals per group (additional data in Figure S7). (B) Limiting dilution analysis of frequency of leukemia initiating cells in secondary *MLL-AF9* transplants. Solid and dashed lines represent the calculated stem cell frequencies and their 95% confidence intervals, respectively. *p* = 6.7 × 10^−8^ for *S222A* versus *MEF2C* (chi-squared test). (C) Western immunoblot analysis for MEF2C and pS222 MEF2C in OCI-AML2 cells lentivirally transduced with wild-type *MEF2C* or dominant negative *MEF2C S222A* transgenes and treated for 48 hours with 600 ng/ml of doxycycline to induce transgene expression. (D) Quantitative analysis of MEF2C and pS222 MEF2C, as measured using quantitative fluorescence immunoblotting, and normalized to actin, demonstrating equal expression of MEF2C and MEF2C S222A protein (\**p* = 0.96, t-test) and significantly reduced abundance of pS222 MEF2C (\*\**p* = 1.1 × 10^−3^ for MEF2C S222A versus MEF2C, t-test). Error bars represent standard deviation of the mean for 3 biological replicates. (E) Growth of human AML cell lines lentivirally transduced with wild-type *MEF2C* or dominant negative *MEF2CS222A* transgenes and treated for 72 hours with 600 ng/ml doxycycline to induce transgene expression.Error bars represent standard deviation of the mean for 3 biological replicates. * and ** *p* = 3.4 × 10^-3^ and 8.4 × 10^−3^ for MEF2C versus MEF2C S222A, respectively (t-test). (F) Kaplan–Meier survival curves of NSG mice transplanted with OCI-AML2 cells transduced with wild-type *MEFC* and dominant negative *MEF2C S222A* transgenes, and treated with doxycycline in chow 3 days following transplantation continuously *in vivo*. *p* = 3.5 × 10^-6^ MEF2C versus MEF2C S222A, log rank test for 10 animals per group.

### MEF2C phosphorylation is required for leukemia maintenance in MLL-rearranged human AML cells

To assess the function of MEF2C S222 phosphorylation in human AML, we analyzed its expression in a panel of human AML cell lines, including both MLL-rearranged and non-rearranged leukemias, and identified OCI-AML2, MOLM-13, MV4-11, and Kasumi-1 cells to have high levels of MEF2C pS222 (Figure S6A). Thus, we used a doxycycline-inducible lentivirus vector to express wild-type and dominant negative MEF2C S222A mutant, and confirmed near-physiologic transgene expression and pS222 levels by Western immunoblotting (Figure 3C-D and S6B). We observed that OCI-AML2 and MOLM-13 AML cell lines with endogenous MEF2C S222 phosphorylation had significantly reduced viability upon the doxycycline-induced expression of MEF2C S222A as compared to wild-type MEF2C (*p* = 8.4 × 10^-3^ and 3.4 × 10^-3^, t-test, respectively, Figure 3E) at least in part due to apoptosis (Figure S6C and S6D). In contrast, HL-60 and U937 AML cells that lack endogenous MEF2C exhibited no significant effects upon expression of wild-type or mutant MEF2C S222A (Figure 3E).

To investigate the function of MEF2C phosphorylation in human leukemias *in vivo*, we next assessed the effects of inhibiting MEF2C S222 phosphorylation by expression of MEF2C S222A in OCI-AML2 cells transplanted orthotopically by tail vein injection in immunodeficient mice (Figure 3F and S6E). Cells were transplanted into NOD-SCID-IL2Rcγnull (NSG) mice, and transgene expression was induced 3 days post-transplant using doxycycline chow. Mice transplanted with OCI-AML2 cells expressing dominant negative MEF2C S222A had significantly prolonged survival and impaired leukemia development as compared to those transplanted with cells expressing wild-type MEF2C or non-transduced control OCI-AML2 cells (*p* = 3.5 × 10^-6^, log-rank test, Figure 3F). These findings indicate that MEF2C S222 phosphorylation is required for the survival of human AML cells *in vitro* and in mouse xenografts *in vivo*.

### Loss of MEF2C S222 phosphorylation impairs MEF2C-mediated transcriptional gene expression program

MEF2C functions as a transcription factor by sequence-specific recognition of MEF2 response element (Yu et al., 1992), through regulated recruitment of transcriptional co-activators and co-repressors via its transactivation (118-473 residues) domain (Kang et al., 2006; Zhu and Gulick, 2004). To test the hypothesis that MEF2C S222 phosphorylation regulates MEF2C transcriptional activity, we expressed wild-type and mutant MEF2C S222A in human embryonic kidney (HEK) 293T cells, and assessed their transactivation of *MEF2* response element-driven firefly luciferase, as compared to CMV promoter-driven *Renilla* luciferase as control (Figure 4A). We confirmed equal MEF2C wild-type and mutant S222A protein expression and reduced pS222 phosphorylation using Western immunoblotting (Figure 4B). Expression of the wild-type MEF2C in HEK293T cells that lack endogenous MEF2C led to an 18-fold increase in MEF2C transcriptional activity as compared to the vector control (Figure 4A). Despite equal levels of wild-type MEF2C and mutant S222A protein expression (Figure 4B), MEF2C transcriptional activity was significantly reduced due to the loss of MEF2C S222 phosphorylation in the S222A mutant (*p* = 4.0 × 10^-2^, t-test, Figure 4A and 4B). This suggests that MEF2C S222 phosphorylation is required for maximal activation of MEF2C-dependent gene transcription.

**Figure 4.**
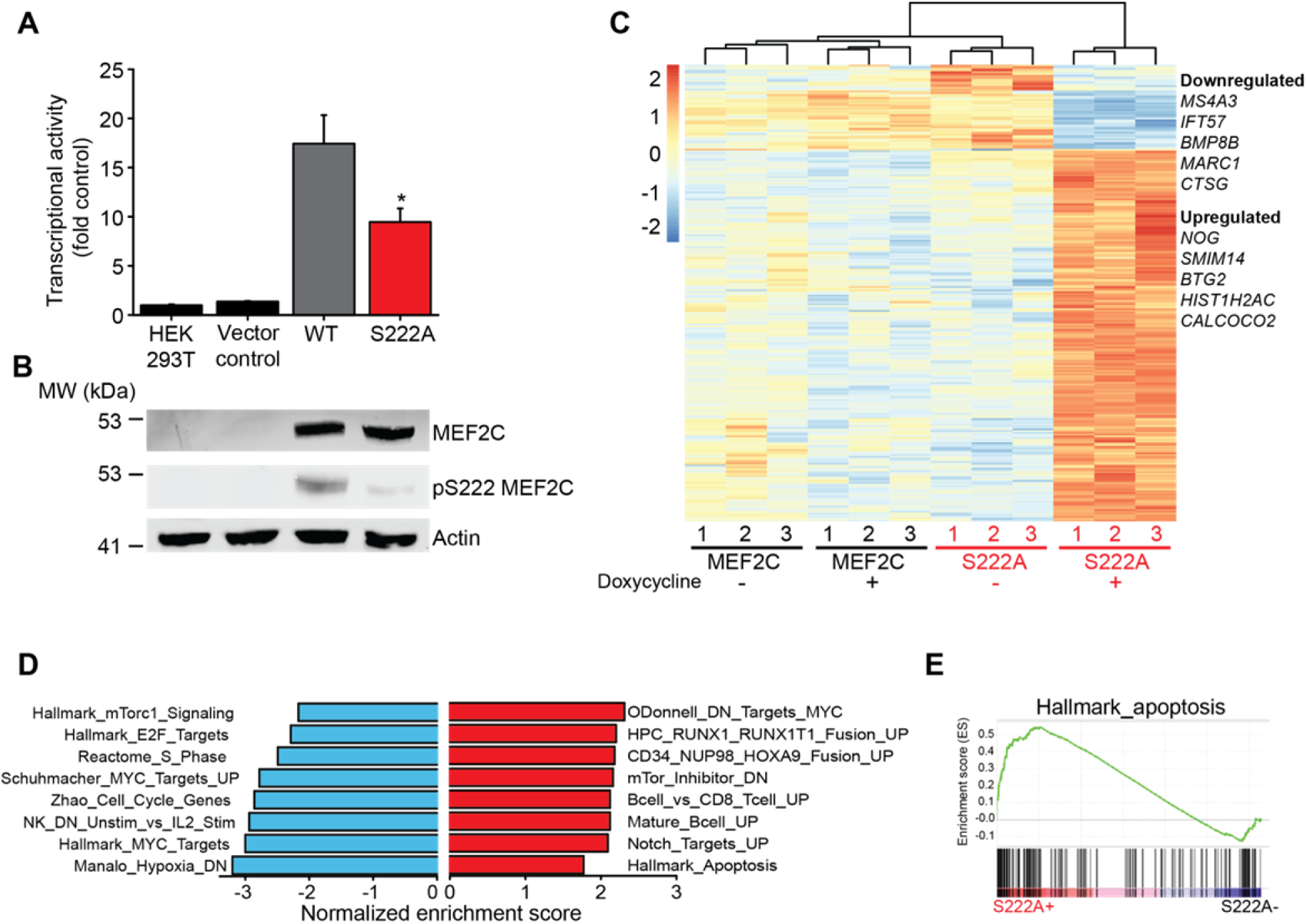
MEF2C phosphorylation is required for MEF2C-mediated gene transactivation. (A) Activity of luciferase transcriptional MEF2 reporter in HEK293T cells lentivirally transduced with wild-type *MEF2C* or mutant *MEF2C S222A*, as compared to vector control. Error bars represent standard deviation of the mean for 3 biological replicates. * *p* = 4.0 × 10^-2^ for MEF2C S222A versus MEF2C, t-test. (B) Western immunoblot analysis for MEF2C and pS222 MEF2C in transcriptional reporter cells, demonstrating equal protein expression of MEF2C transgenes, and reduced pS222 in MEF2C S222A transduced cells. (C) Hierarchical clustering of gene expression of the most differentially expressed genes in OCI-AML2 cells transduced with wild-type *MEF2C* or dominant-negative *MEF2C S222A* transgenes (-), and treated with 600 ng/ml doxycycline for 48 hours to induce transgene expression (+). Three biological replicates are shown, as indicated. Blue-to-red color gradient indicates relative gene expression. (D) Gene set enrichment analysis (GSEA) of significantly upregulated and downregulated gene sets. (E). GSEA illustrates enrichment of apoptotic genes in MEF2C S222A expressing cells. Normalized enrichment score = 1.76 and false discovery rate q = 5.2 × 10^-3^.

To elucidate the gene expression program controlled by MEF2C S222 phosphorylation in AML cell survival, we analyzed gene expression profiles of OCI-AML2 cells expressing wild-type as compared to dominant negative MEF2C S222A using RNA sequencing (Figure 4C). Consistent with the near-physiologic expression of MEF2C transgenes (Figure 3C), we found essentially no significant differences in gene expression profiles of cells expressing wild-type MEF2C upon doxycycline-induced transgene expression (Figure 4C). In contrast, we observed a significant change in gene expression of cells upon doxycycline-induced expression of MEF2C S222A, as compared to both un-induced cells and cells expressing wild-type MEF2C (Figure 4C). In particular, we found 276 genes that were significantly altered in expression in cells expressing MEF2C S222A as compared to un-induced cells (Figure 4C, Data S2). Using quantitative reverse transcriptase polymerase chain reaction (qRT-PCR), we found that none of the canonical MEF2C target genes, such as *Nr4a1/Nur77*, *Hdac7* and *Jun*, were significantly changed in expression upon expression of MEF2C S222A (Figure S5I-K). Instead, gene set enrichment analyses revealed that loss of MEF2C S222 phosphorylation led to downregulation of distinct gene expression programs, including E2F and MYC target genes (Figure 4D, Data S3-4). In addition, we observed altered expression of genes regulating apoptosis (Figure 4D and 4E, Data S3-4), consistent with the phenotypic induction of apoptosis upon the blockade of MEF2C S222 phosphorylation in mouse and human leukemia cells (Figures S5D and S6C–D).

### MARK-mediated phosphorylation of MEF2C S222 confers susceptibility to MARK inhibitors in AML

The functional requirements of MEF2C S222 phosphorylation for leukemia initiation and maintenance, combined with its dispensability for normal hematopoiesis, suggest that blockade of MEF2C phosphorylation may be exploited for anti-leukemic therapy. Given that direct inhibition of transcription factors is pharmacologically challenging, we reasoned that inhibition of the upstream kinases phosphorylating MEF2C at S222 could downregulate MEF2C signaling to impair AML cell survival. To identify candidate kinases that can phosphorylate S222 of MEF2C, we screened a library of 172 recombinant human serine kinases in their ability to phosphorylate a model substrate in the presence of synthetic pS222-containing peptide as a competitive product inhibitor (Figure 5A, Data S5). We found that only five protein kinases scored significantly in this screen, of which four belonged to a closely related microtubule-associated protein/microtubule affinity regulating kinase (MARK) 1, 2, 3, 4 family (Figure 5A). Analysis of primary AML specimens revealed that *MARK3* and *MARK2* genes are highly expressed in AML (Figure S7A). We validated the specific activity of recombinant MARK3 with synthetic MEF2C pS222 peptide, as compared to the closely related CAMK1α kinase (Figure 5B). These results indicate that MARK3 and related MARK kinase family members can specifically phosphorylate S222 MEF2C *in vitro*.

**Figure 5.**
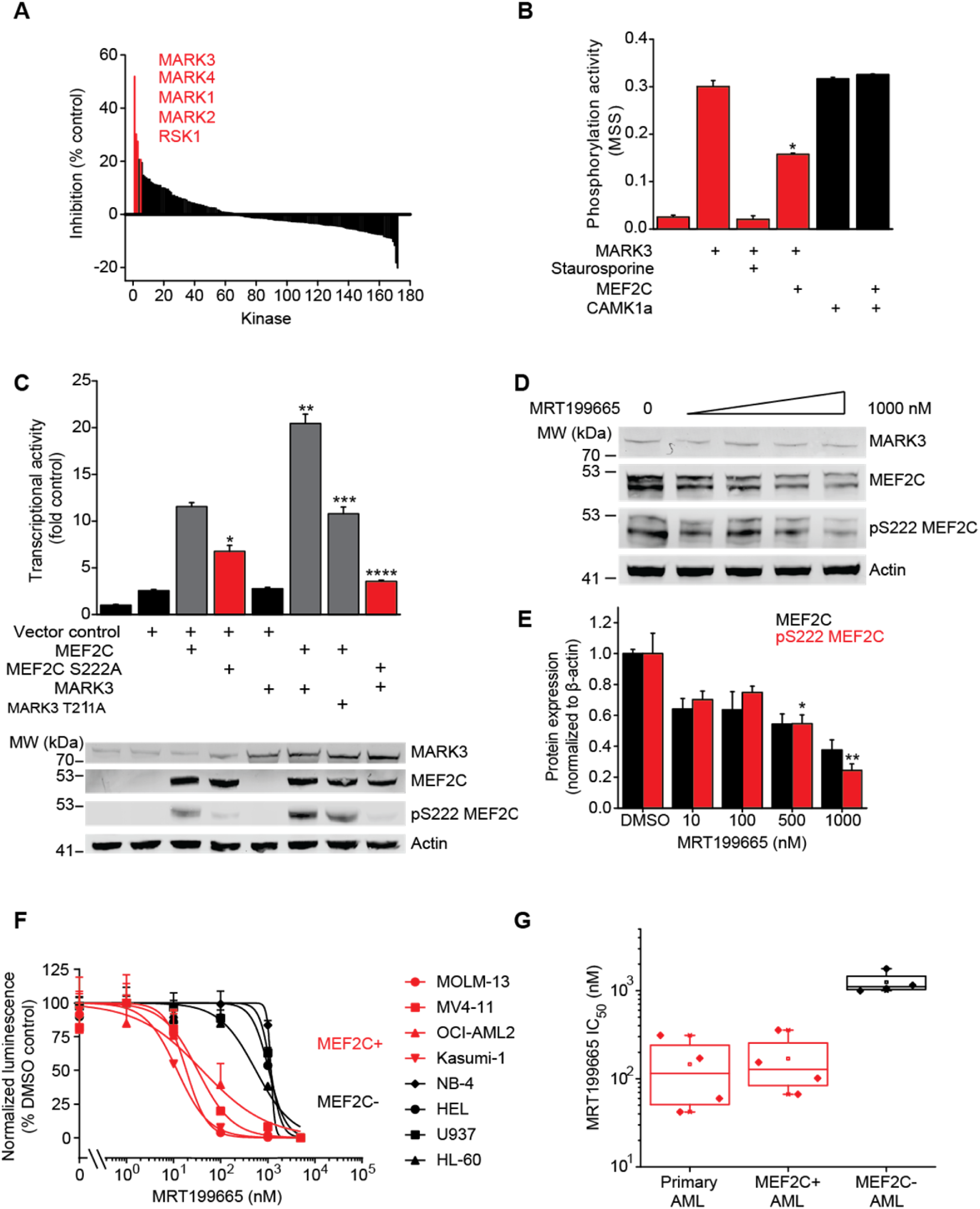
Chemical inhibition of MARK-induced MEF2C phosphorylation exhibits selective toxicity against MEF2C-activated human AML cells. (A) Recombinant screen for serine kinases that phosphorylate MEF2C S222, as assayed by significant pS222 MEF2C product inhibition marked in red. (B) Phosphorylation activity of recombinant MARK3 (red) as compared to control CAMK1α (black) on model substrate as product-inhibited by synthetic pS222 MEF2C peptide. Staurosporine serves as positive control. Error bars represent standard deviation of the mean for 3 biological replicates. * *p* = 4.1 × 10^-5^ for MARK3 activity with and without MEF2C, t-test. (C) Activity of luciferase transcriptional MEF2 reporter in HEK293T cells lentivirally transduced with MEF2C, MARK3 or their mutants, as indicated. Error bars represent standard deviations of the mean for 3 biological replicates. *, **, *** and **** *p*= 2.3 × 10^-4^, 3.4 × 10^-5^, 5.0 × 10^-5^, and 1.5 × 10^-7^ for MEF2C S222A; vector versus MEF2C;vector, MEF2C;MARK3 versus MEF2C;vector, MEF2C;MARK3 T211A versus MEF2C;MARK3, and MEF2C S222A;MARK3 versus MEF2C;MARK3, respectively (t-test). Below, Western immunoblot analysis demonstrating equal protein expression of MEF2C and MARK3 transgenes, with reduced S222 phosphorylation by expression of MEF2C S222A and MARK3 T2111A mutants. (D) Western immunoblot for MEF2C, pS222 MEF2C and MARK3 in OCI-AML2 cells treated with increasing concentrations of MRT199665 for 12 hours. (E) Quantitative analysis of MEF2C and pS222 MEF2C abundance, as measured using quantitative fluorescence immunoassays, and normalized to actin, demonstrating significant reduction of MEF2C phosphorylation by MRT199665 as compared to DMSO control (*p* = 5.5 × 10^-3^ and 3.4 × 10^-4^ for 500 and 1000 nM, respectively (t-test). Error bars represent standard deviation of the mean for 2 biological replicates. (F) Growth of human AML cell lines at 48 hours as a function of MRT199665 concentration for MEF2C-expressing cells (red) as compared to those that lack MEF2C (black), as indicated. Error bars represent standard deviation of the mean for 3 biological replicates. (G) MRT199665 *IC*_*50*_ values for primary patient AML specimens and human AML cell lines with (red) and without (black) MEF2C activation upon 48 hours of drug treatment *in vitro*. Box plots represent mean and quartile, with whiskers denoting maximum and minimum values (additional data in Figure S11).

To evaluate whether MARK3 could phosphorylate MEF2C S222 and regulate its transcriptional activity in cells, we co-expressed MEF2C and MARK3 in HEK293T cells and assessed MEF2C phosphorylation and transcriptional activity by Western immunoblotting and transcriptional reporter assays, respectively (Figure 5C). We found that MEF2C transcriptional activity was significantly increased upon co-expression of MEF2C and MARK3 as compared to cells expressing MEF2C alone (*p* = 3.4 × 10^-5^, t-test, Figure 5C). This increased transcriptional reporter activity was associated with increased levels of pS222, which was blocked by the co-expression of the enzymatically impaired MARK3 T211A activation loop mutant (Nesic et al., 2010; Timm et al., 2008) (*p* = 5.0 × 10^-5^, Figure 5C). Importantly, expression of MARK3 failed to activate the transcriptional activity or induce S222 phosphorylation of the MEF2C S222A mutant (Figure 5C). Taken together, these data indicate that MARK kinase signaling can regulate MEF2C transcriptional activity specifically in a S222 phosphorylation-dependent manner.

To test the hypothesis that inhibition of MARK-mediated phosphorylation of MEF2C may have anti-leukemic efficacy, we identified the ATP-competitive kinase inhibitor MRT199665 which exhibits high selectivity and potency against MARK kinases as compared to other structurally AMP kinases (Clark et al., 2012). Treatment of OCI-AML2 cells with increasing concentrations of MRT199665 led to a dose-dependent reduction in total and pS222 MEF2C (Figure 5D), causing more than 40% reduction in MEF2C phosphorylation at 10 nM as compared to untreated cells, as assessed by quantitative fluorescent Western immunoblotting (Figure 5E). We observed that human AML cell lines with endogenous MEF2C phosphorylation (OCI-AML2, MV4-11, MOLM-13 and Kasumi-1) were more sensitive to MRT199665 as compared to cell lines lacking MEF2C (NB-4, HEL, HL-60 and U937) with mean 50% inhibitory concentrations (*IC*_*50*_) of 26 ± 13 versus 990 ± 29 nM, respectively (*p* = 5.6 × 10^-5^, Figure 5F), and displayed reduced leukemia growth in clonogenic assays (Figure S7B). Likewise, we found that primary patient AML specimens with MEF2C activation treated in short-term cultures *ex vivo* exhibited enhanced susceptibility and apoptosis in response to MRT199665 treatment as compared to AML cells lacking MEF2C expression (*IC*_*50*_ = 150 ± 120 versus 1300 ± 360 nM, respectively, *p* = 1.8 × 10^-5^, Figures 5G and S7C–F). Thus, activation of MEF2C in human AML confers susceptibility to MARK kinase inhibition.

### MEF2C phosphorylation is required for chemotherapy resistance in AML

Since MEF2C phosphorylation was specifically observed in diagnostic AML specimens in patients with failure of induction therapy (Figure 1), we hypothesized that MEF2C S222 phosphorylation induces chemotherapy resistance. In agreement with this prediction, OCI-AML2 cells induced to express dominant negative MEF2C S222A were significantly more sensitive to both cytarabine and doxorubicin treatment as compared to cells expressing wild-type MEF2C or untransduced OCI-AML2 cells (*p* = 8.8 × 10^-8^ and *p* = 3.4 × 10^-9^, respectively, chi-squared test, Figure 6A and 6B). We found that human AML cell lines with endogenous MEF2C phosphorylation (OCI-AML2, MV4-11, MOLM-13 and Kasumi-1) were significantly sensitized to cytarabine in the presence of 10 nM MRT199665, as compared to cell lines lacking MEF2C (NB-4, HEL, HL-60 and U937) that remained unaffected by MRT199665 (*p* = 0.024, paired t-test, Figure 6C). Similarly, primary patient AML specimens with MEF2C activation exhibited enhanced susceptibility to cytarabine treatment in the presence of MRT199665 *in vitro* (*IC*_*50*_ = 58 ± 76 versus 230 ± 280 nM in the presence versus absence of MRT199665, *p* = 0.036, t-test, Figure 6D). Taken together, these results indicate that MEF2C phosphorylation is required for AML chemotherapy resistance, which can be blocked by MARK kinase inhibition.

**Figure 6.**
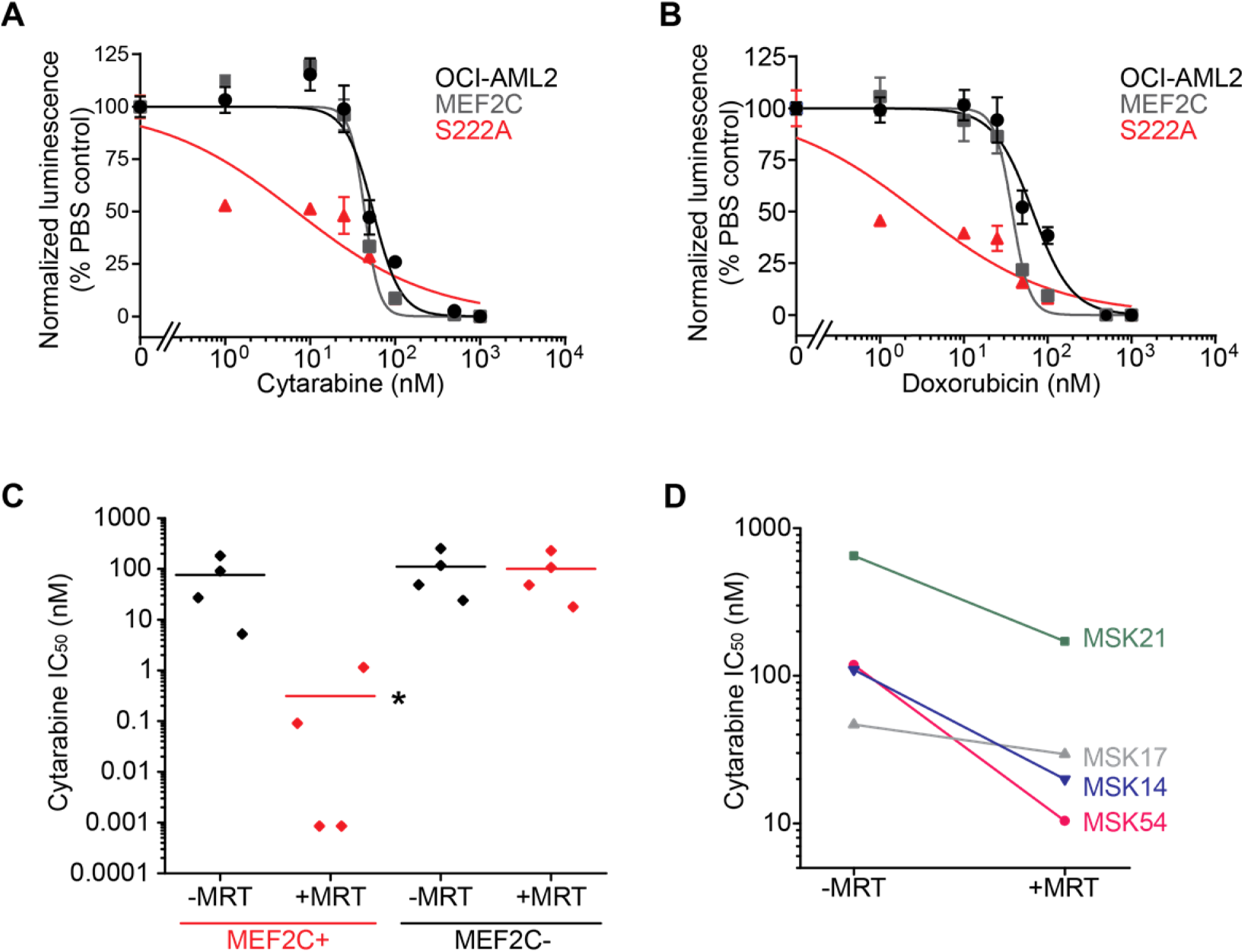
MARK kinase inhibition overcomes chemotherapy resistance of MEF2C-activated AML cell lines and patient cells. (A) Growth of OCI-AML2 cells lentivirally transduced with wild-type *MEF2C* or dominant negative *MEF2C S222A* transgenes and treated with 600 ng/ml doxycycline to induce transgene expression and increasing concentrations of cytarabine (A) and doxorubicin (B) for 72 hours. Error bars represent standard deviation of the mean for 3 biological replicates. *p* = 8.8 × 10^-8^ and 3.4 × 10^-9^ for MEF2C versus MEF2C S222A by nonlinear regression for cytarabine and doxorubicin, respectively. (C) Cytarabine *IC*_*50*_ values for human AML cell lines with MEF2C-activation (MEF2C+) as compared to those lacking MEF2C (MEF2C-) after 48 hours of drug treatment in the absence (-MRT) or presence of 100 nM (+MRT) MRT199665. Each data point represents the mean of biological triplicates of an individual sample. * *p* = 0.024 for +MRT versus– MRT for MEF2C-activated cells by paired t-test. (D) Cytarabine *IC*_*50*_ values for primary patient AML specimens with MEF2C after 48 hours of drug treatment in the absence (-MRT) or presence of 100 nM (+MRT) MRT199665 *in vitro*. Each data point represents the mean of biological triplicates of an individual sample. * *p* = 0.036 for +MRT verses–MRT by paired t-test.

## DISCUSSION

Our current findings indicate that MEF2C S222 phosphorylation is a specific marker of primary chemotherapy resistance and failure of induction therapy in patients with both cytogenetically normal and chromosomally-rearranged acute myeloid leukemias (Figure 1). Activity of S222-phosphorylated MEF2C appears to be aberrant, insofar as mice genetically engineered to block its phosphorylation exhibit normal steady-state and stress hematopoiesis, but are resistant to leukemogenesis induced by the *MLL-AF9* oncogene *in vivo* (Figure 2). At least in part, this effect is due to the requirement for MEF2C phosphorylation for the survival of leukemia stem or initiating cells (Figure 3), and enhanced transcriptional activity and induction of gene expression programs regulating cell survival and apoptosis (Figure 4). Finally, MARK kinases can specifically phosphorylate MEF2C, potentiating its transcriptional activity (Figure 5), and MARK-selective kinase inhibition can overcome chemotherapy resistance of MEF2C-activated human AML cell lines and patient leukemias (Figure 6).

Resistance to chemotherapy remains the major barrier to improving clinical outcomes for patients with AML, yet our current concepts of chemoresistance lack sufficient explanatory power. For example, inactivation of TP53 and overexpression of xenobiotic transporters have been found to cause chemoresistance, but the majority of patients with primary refractory or relapsed AML lack identifiable mutations of TP53 or known components of TP53-dependent DNA damage response, and do not exhibit xenobiotic transporter overexpression (Cancer Genome Atlas Research, 2013; Papaemmanuil et al., 2016; Schuback et al., 2013). Likewise, we do not yet understand how recently identified groups of mutations associated with poor prognosis cause chemotherapy resistance (Papaemmanuil et al., 2016). Recently, impaired susceptibility to mitochondrial apoptosis has been associated with chemoresistance in AML (Pan et al., 2014; Pierceall et al., 2013). However, the molecular mechanisms responsible for these differences also remain poorly understood. Our findings identify kinase-dependent dysregulation of transcription factor control as a determinant of therapy response in AML, at least in part mediated by aberrant survival and apoptosis resistance of leukemia stem cells (Figure 7). This suggests that varied mechanisms of chemotherapy resistance and disease relapse in patients may be ultimately caused by aberrant gene expression programming of privileged leukemia cell subsets that provide a chemoresistant disease reservoir (Goardon et al., 2011).

**Figure 7.**
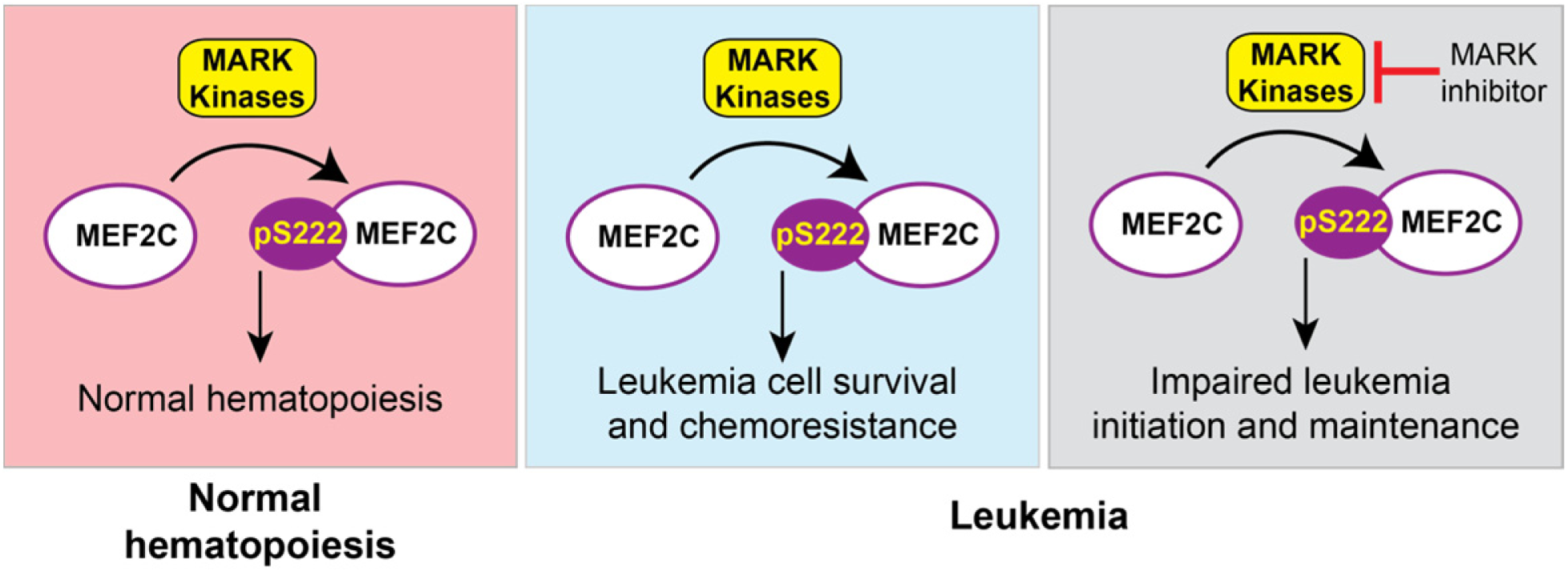
Model of signaling-dependent dysregulation of MEF2C in AML and chemotherapy resistance. MEF2C S222 phosphorylation is regulated by MARK kinases and is dispensable for normal hematopoiesis, but is required for enhanced leukemia stem cell survival and chemotherapy resistance. This differential functional dependency can be exploited for therapy and blockade of chemotherapy resistance using selective MARK kinase inhibition.

The observation of MEF2C S222 phosphorylation in genetically diverse AML subtypes, including cytogenetically normal and MLL-rearranged leukemias, raises the possibility that common gene expression programs and regulatory mechanisms may be engaged by distinct genetic and molecular classes of AML, such as for example distinct groups of mutations and/or their induction in specific leukemia-initiating cell populations. Likewise, recurrent mutations of *MEF2B*, *MEF2C* and *MEF2D* in refractory lymphoid cancers (Gu et al., 2016; Pon et al., 2015; Ying et al., 2013), suggest that MEF2 family members may regulate essential survival or homeostatic mechanisms in hematopoietic cells which cause therapy resistance in myeloid and lymphoid malignancies. However, whereas mutational activation of *MEF2C* in refractory T-cell acute lymphoblastic leukemias appears to inhibit apoptosis via NR4A1/NUR77-mediated effects on BCL2 (Nagel et al., 2008), apoptosis resistance induced by MEF2C phosphorylation in AML appears to involve distinct mechanisms (Figure 4). We observe that MEF2C is both phosphorylated and highly expressed, and its leukemogenic activities may therefore be due to both its high abundance and S222 phosphorylation. Aberrant MEF2C S222 phosphorylation may affect its activity by i) recruitment of new co-activators or co-repressors, ii) altered dimerization with endogenous MEF2 family members, and/or iii) altered target gene localization due to secondary changes in the post-translational modifications and sequence binding preferences of its DNA binding domain (Barneda-Zahonero et al., 2013; Kim et al., 2005; Lu et al., 2000; Mao et al., 1999). Since our functional studies were largely based on *MLL-AF9*-rearranged leukemias, it is possible that MEF2C phosphorylation may involve distinct molecular mechanisms in different leukemia subtypes. It is also possible that the marked phenotype of non-phosphorylatable MEF2C mutants may be due to dominant negative effects on other MEF2 family members. Similarly, while MARK3 can specifically phosphorylate MEF2C S222, which is inhibited by MRT199665, additional serine kinases which were not included in our screen, such as mTOR for example, may induce MEF2C phosphorylation in distinct types of AML.

The MARK family was originally discovered based on its functions in controlling cell polarity and microtubule dynamics as part of the basolateral polarity complex (Guo and Kemphues, 1995; Shulman et al., 2000), but their functions in hematopoietic cells are not yet defined (Hurov et al., 2007). Our finding that MARKs can regulate MEF2C suggest that the basolateral polarity complex may have distinct functions in normal hematopoiesis which may be dysregulated in leukemia cells. Similarly, the mechanisms of MARK3-mediated phosphorylation of MEF2C in chemotherapy-resistant AML remain to be defined. For example, MARK kinases can be phosphorylated and activated by GSK-3β (Kosuga et al., 2005), which can in turn by activated by integrin signaling, including reports of its activation in AML (Miller et al., 2013). Thus, activation of MARK signaling and MEF2C phosphorylation may be linked to autocrine or paracrine signaling by AML cells and the bone marrow niche (Kentsis et al., 2012; Zahreddine et al., 2014; Zhou et al., 2016).

Finally, the lack of apparent phenotypes in knock-in mutant mice homozygous for *Mef2c*^*S222A/S222A*^ or *Mef2c*^*S222D/S222D*^ suggests that phosphorylation of MEF2C S222 is dispensable for normal development, establishing a compelling therapeutic window and substantiating its therapeutic targeting. However, additional studies of MEF2C phosphorylation in specific physiologic states (Herglotz et al., 2016; Wang et al., 2016), and further preclinical development of selective MARK inhibitors will be needed to advance their use as MEF2C-targeted therapeutics. Future functional profiling studies of large AML cohorts should establish the prevalence of MEF2C phosphorylation and other mechanisms of primary chemotherapy resistance.

## EXPERIMENTAL PROCEDURES

### Collection of patient samples

Written informed consent and approval by the Institutional Review Boards of participating institutions was obtained in accordance with the Declaration of Helsinki for all subjects. Primary chemotherapy resistance was defined based on the presence of at least 5% of abnormal blasts by morphologic and immunophenotypic assessment of bone marrow aspirates obtained after two cycles of induction chemotherapy, as assessed by the respective institutional or central pathologic reviews. Specimens were collected and leukemia cells purified as previously described (Brown et al, 2016). Mononuclear cells were purified using Ficoll gradient centrifugation, and leukemia cells were purified by negative immunomagnetic selection against CD3, CD14, CD19 and CD235a, based on the absence of their expression by the majority of AML specimens.

### Production and purification of pS222 MEF2C antibody

A phospho-specific antibody against MEF2C S222 (RefSeq ID: NM_002397.4) was generated by PhosphoSolutions (Catalog p1208-222, RRID:AB_2572427; PhosphoSolutions, Aurora, CO, USA). Rabbits were immunized with a short synthetic phospho-peptide corresponding to amino acids surrounding S222 of human MEF2C, conjugated with Keyhole limpet hemocyanin (KLH). Serum isolated from peripheral blood of immunized rabbits was screened for phospho-specificity using enzyme-linked immunosorbent assays (ELISA). ELISA-positive sera were pooled and sequentially affinity-purified using both the phospho-peptide and non-phospho peptide columns to isolate the affinity-purified pS222 MEF2C antibody.

### Mouse studies

All mouse studies were conducted with approval from the Memorial Sloan Kettering Cancer Center Institutional Animal Care and Use Committee. C57BL/6J, NOD-Scid IL2Rγ-null (NSG) and B6.SJL-Ptprca Pepcb/BoyJ mice were all obtained from Jackson Labs (Bar Harbor, ME, USA). Generation of knock-in *Mef2c*^*S222A*^ and *Mef2c*^*S222D*^ mutant mice using CRISPR/Cas9 genome editing and homologous recombination is described in the Supplemental Experimental Procedures. For all experiments using *Mef2c* knock-in mutant animals, wild-type litter mates were used as controls.

### Phosphoproteomics screen

Phosphoproteomic profiling was performed as described (Ficarro et al., 2011). Briefly, purified leukemia cells were lysed using guanidine hydrochloride, and proteins were reduced with dithiothreitol, alkylated with iodoacetamide, and digested using trypsin. Tryptic peptides were purified by solid phase extraction, and purified peptides were labeled using iTRAQ reagent. Phosphopeptides were enriched using iron affinity chromatography, and separated using three-dimensional RP-SAX-RP chromatography coupled to nanoelectrospray ion source. Spectra were recorded using Orbitrap Velos mass spectrometer (ThermoFisher Scientific, San Jose, CA) in data dependent mode. Data files were analyzed using multiplierz (Parikh et al., 2009).

### Recombinant kinase screen

The protein kinase screen was performed using a recombinant serine kinase library as previously described (Kitagawa et al., 2012) and detailed in the Supplemental Experimental Procedures.

### Data and statistical analysis

Statistical significance was determined using two-tailed Student’s t-test for continuous variables, Pearson’s chi-squared test for limiting dilution assays, and log-rank test for survival analysis.

Detailed description of the methods is provided as part of the Supplemental Information.

## AUTHOR CONTRIBUTIONS

Methodology, F.C.B., P.C., C.R., S.T., W.M., P.R., N.G., J.M. and A.K., Software, R.P.K., A.K.; Validation, F.C.B., E.S., P.C., P.R., N.G. and A.K., Formal Analysis, F.C.B., R.P.K., B.S. and A.K.; Investigation, F.C.B., E.S., P.C., S.T., C.O.D., P.R., N.G. and A.K.; Resources, G.P., A.V. K. and S.A.A,; Writing – Original Draft, F.C.B. and A.K.; Writing – Review and Editing, F.C.B. and A.K.; Visualization, F.C.B., E.S., R.P.K., J.M. and A.K. Conceptualization, Supervision, Funding Acquisition, A.K.

## ACKNOWLEDGEMENTS

We are grateful to Alejandro Gutierrez, Marc Mansour, Leo Wang, Michael Kharas, Martin Tallman, and Ross Levine for critical discussions. We thank John Schwarz and Hanna Mikkola for Mef2c conditional mice, Philip Cohen for MRT199665, and Antoine Gruet, Matthew Witkin and Yang Li for technical assistance. This work was supported by the NIH R21 CA188881, R01 CA204396, P30 CA008748, Burroughs Wellcome Fund, Josie Robertson Investigator Program, Rita Allen Foundation, Alex’s Lemonade Stand Foundation, American Society of Hematology, and Gabrielle’s Angel Foundation. A.K. is the Damon Runyon-Richard Lumsden Foundation Clinical Investigator.

---

**Figure S1.**
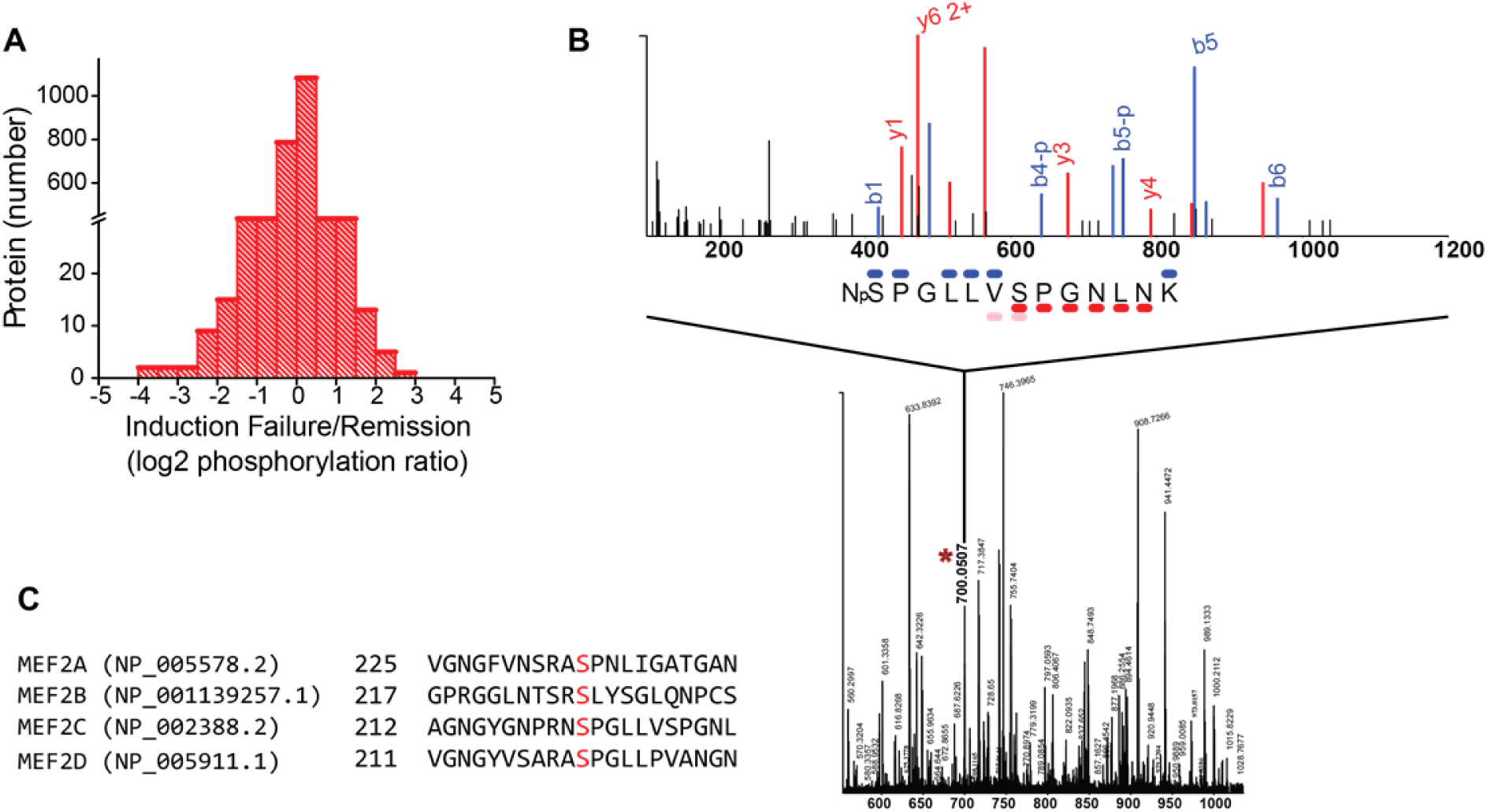
Related to Figure 1. MEF2C phosphorylation is identified in a phosphoproteomic screen for primary chemoresistance. (A) Distribution of differentially expressed phosphorylated proteins in diagnostic AML specimens with primary chemotherapy resistance and induction failure as compared to complete induction remission. (B) MS1 plots of pS222 MEF2C. (C) Protein sequence alignment of the MEF2 isoforms at the sequence surrounding S222 of MEF2C (shown in red). Isoform 1 has been used for each alignment.

**Figure S2.**
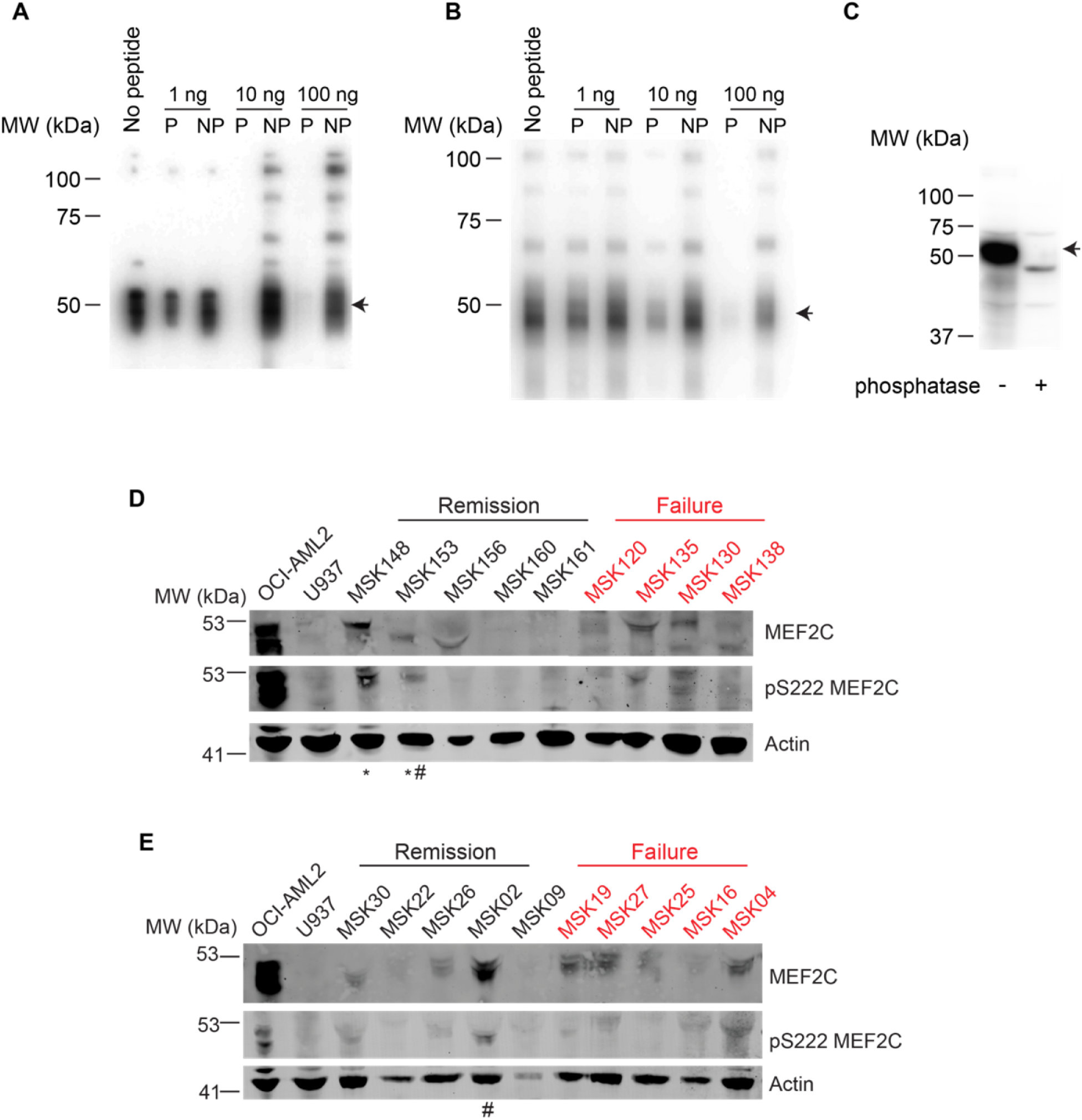
Related to Figure 1. pS222 MEF2C is associated with chemoresistance in primary AML. (A)Peptide competition Western immunoblot of OCI-AML2 cells for pS222 MEF2C. P = phosphorylated MEF2C peptide at S222; NP = non-phosphorylated MEF2C peptide at S222. (B) Peptide competition Western immunoblot for pS222 MEF2C of HEK293T cells lentivirally transduced with wild-type MEF2C. (C) Lysates prepared from HEK293T cells lentivirally transduced with wild-type MEF2C were treated with (+) or without (−) λ phosphatase (800 units) and immunoblotted for pS222 MEF2C. Arrows indicate pS222 MEF2C. Representative Western Immunoblot analysis for MEF2C and pS222 MEF2C in pediatric (B) and adult (C) age, disease and therapy-matched patient AML specimens with induction failure and complete remission. The human AML cell lines OCI–AML2 and U937 were used for positive and negative controls for MEF2C expression, respectively. ^#^ and * denotes specimens from patients who achieved complete remission but experienced AML relapse, and those with *MLL/KMT2A* cryptic rearrangement.

**Figure S3.**
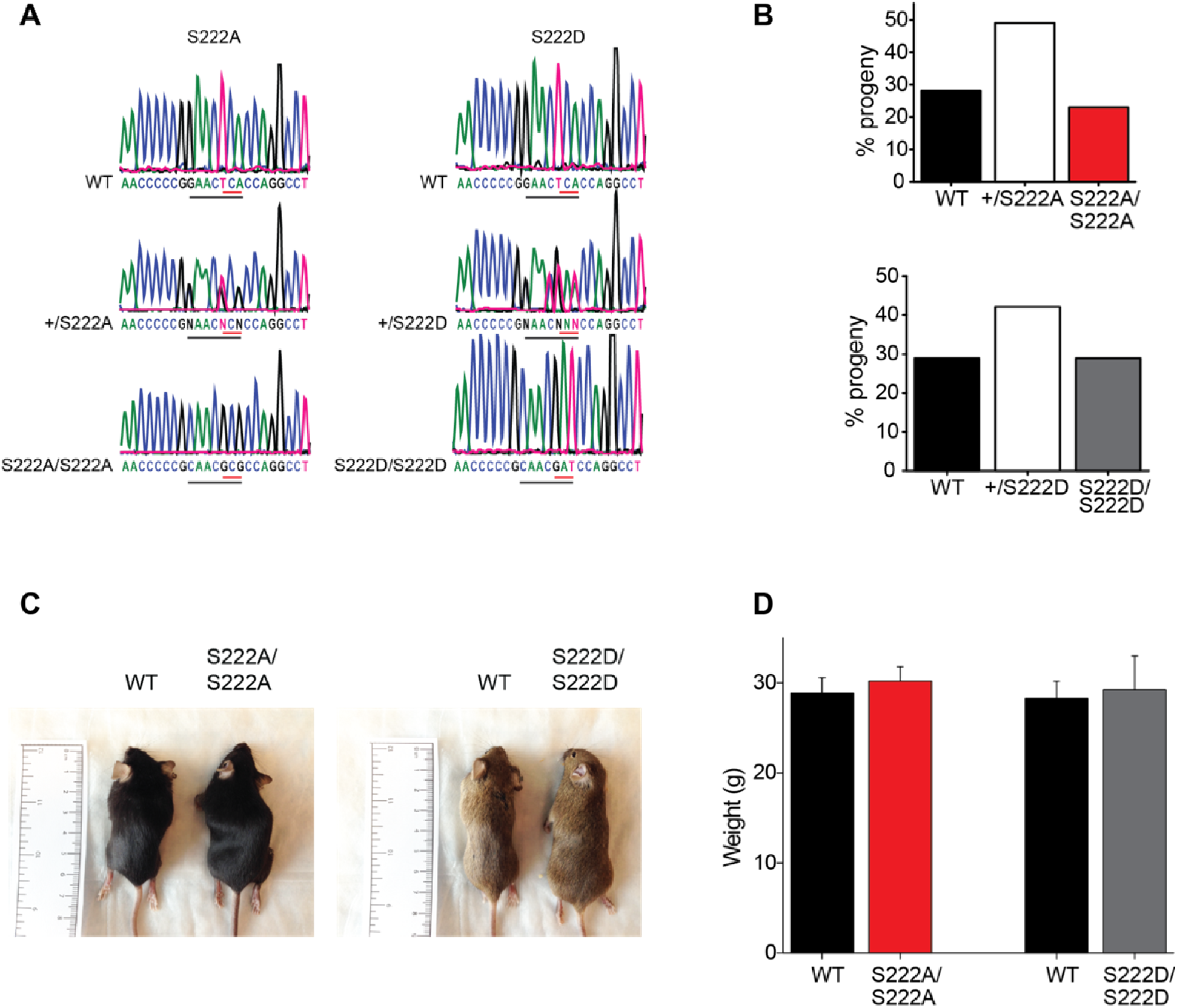
Related to Figure 2. *Mef2c*^*S222A/S222A*^ and *Mef2c*^*S222D/S222D*^ mice develop normally. (A) Sequencing electropherograms of tail genomic DNA from *Mef2c*^*S222A*^ and *Mef2c*^*S222D*^ mice at the *Mef2c* locus. CRISPR/Cas9-induced genome editing introduces c. TCA>GCG (S222A) and c. TCA>GAT (S222D) mutations, and restriction enzyme sequencing sites, as underlined red and black, respectively. (B) Mendelian ratios from heterozygous intercrosses for *Mef2c*^*S222A*^ and *Mef2c*^*S222D*^ mice. Error bars represent standard deviation of the mean. Body size (C) and weight (D) is preserved in *Mef2c*^*S222A/S222A*^ and *Mef2c*^*S222D/S222D*^ mice compared to wild-type litter mate controls. Error bars represent standard deviation of the mean.

**Figure S4.**
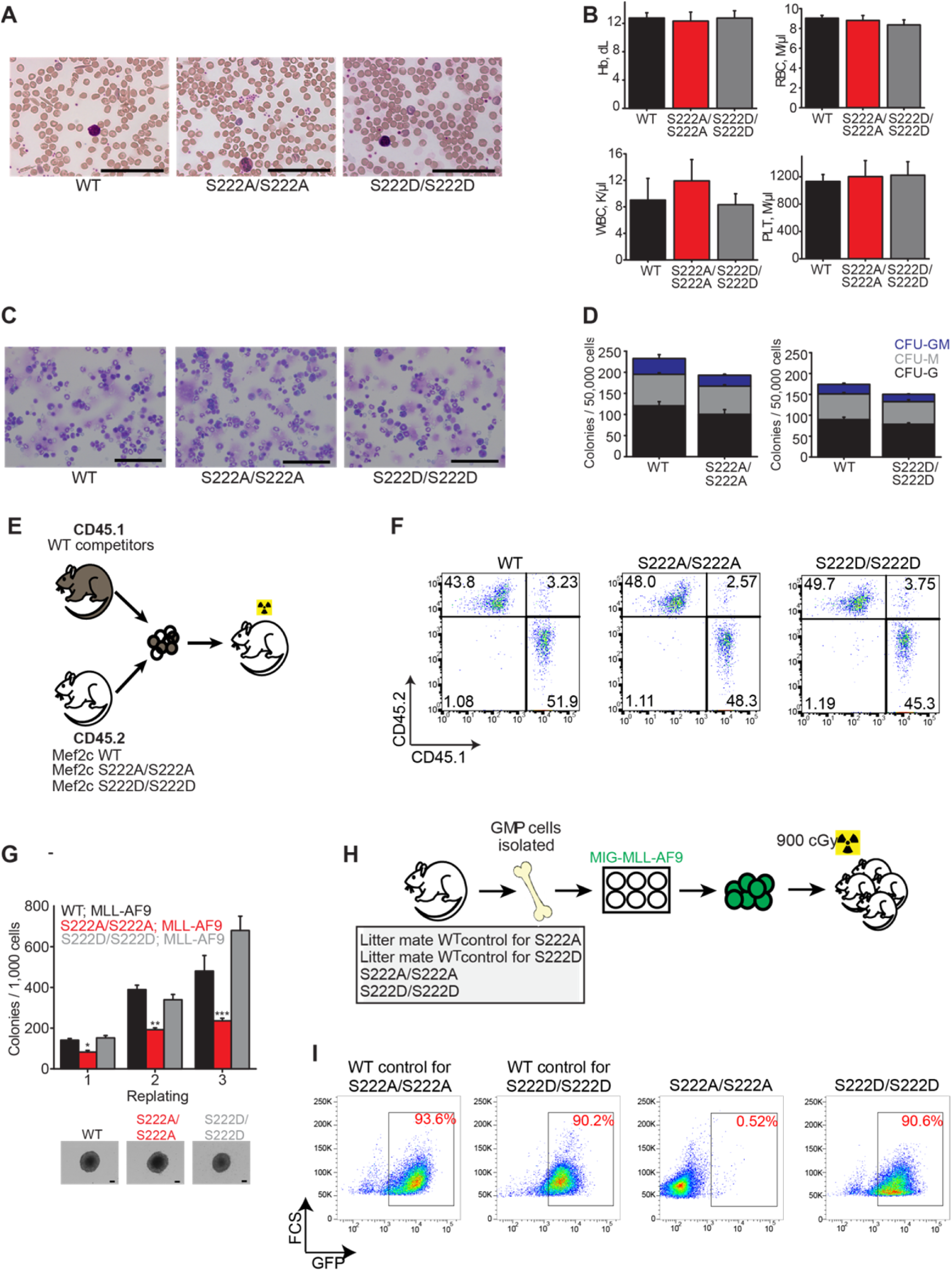
Related to Figure 2. Mef2c phosphorylation is dispensable for normal hematopoiesis but required for the development of MLL-AF9 leukemia *in vivo*. (A) Representative micrographs of peripheral blood smears from *Mef2c*^*S222A/S222A*^ and *Mef2c*^*S222D/S222D*^ mice. Scale bar denotes 50 µm. (B) Enumeration of hemoglobin (Hb), red blood cells (RBC), white blood cells (WBC) and platelets (PLT) in peripheral blood from *Mef2c*^*S222A/S222A*^ and *Mef2c*^*S222D/S222D*^ mice compared to wild-type litter mates. Error bars represent standard deviation of the mean of six animals per genotype. (C) Representative images of Wright-Giemsa–stained cytospin preparations of bone marrow cells from *Mef2c*^*S222A/S222A*^ and *Mef2c*^*S222D/S222D*^ mice. Scale bar denotes 50 µm. (D) Colony formation of GMP cells from *Mef2c*^*S222A/S222A*^ and *Mef2c*^*S222D/S222D*^ mice. Error bars represent standard deviation of the mean of 3 biological replicates. Abbreviations: colony-forming units granulocyte/macrophage (CFU-GM); colony-forming units granulocyte (CFU-G); colony–forming units granulocyte/macrophage (CFU-M) (E) Schematic of competitive transplant. (F) Representative FACS plot of peripheral blood mononuclear cell CD45.2 mutant and CD45.1 wilt–type cells 4 weeks following competitive transplantation. (G) Serial replating clonogenic efficiencies of bone marrow granulocyte-macrophage progenitor cells from an independent mouse strain compared to Figure 2G for *Mef2c*^*S222A/S222A*^, *Mef2c*^*S222D/S222D*^ and wild-type litter mates transduced with *MLL–AF9*. Error bars represent standard deviation of the mean of 3 biological replicates. Below, representative micrographs of colonies at day 7. Scale bar denotes 100 µm. *, ** and *p* = 8.7 × 10^−4^, 1.2 × 10^−4^and 6.2 × 10^−4^of *WT; MLL-AF9* versus *S222A/S222A; MLL-AF9,* t–test. (H)Schematic for *in vivo Mef2c*^*S222A*^ and *Mef2c*^*S222D*^*; MLL–AF9* experiments. (I) Representative FACS plot analysis of bone marrow cells collected from primary recipient animals transplanted as shown in (H). Wild–type (WT) and *Mef2c*^*S222D/S222D*^ cells were collected from moribund animals. *Mef2c*^*S222A/S222A*^ cells were collected from bone marrow aspirates 170 days after transplant.

**Figure S5.**
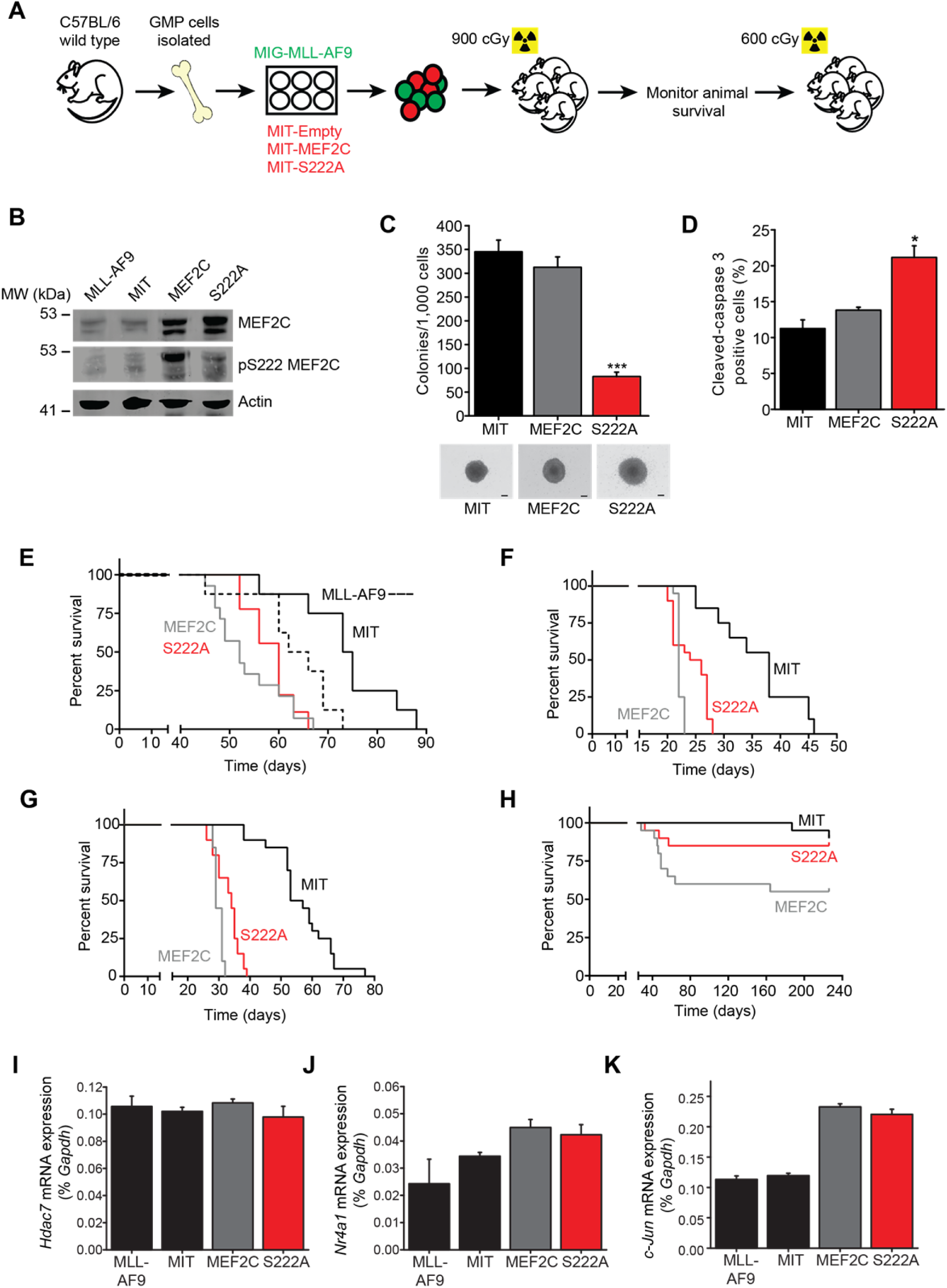
Related to Figure 3. MEF2C phosphorylation is required for leukemia stem cell survival and maintenance in MLL-rearranged leukemia. (A) Schematic for *in vivo MLL-AF9* and *MEF2C* initiation/maintenance experiments. (B) Western immunoblot analysis for MEF2C and pS222 MEF2C in primary leukemia cells shown in (A). (C) Colony formation of primary leukemia cells shown in (A). Below, representative micrographs of colonies at day 7. *** *p* = 5.1 × 10^-5^ S222A vs MEF2C, t-test. Error bars represent standard deviation of the mean of 3 biological replicates. Scale bar denotes 100 µm. (D) Cleaved-caspase 3 analysis of primary leukemia cells shown in (A). Error bars represent standard deviation of the mean of 3 biological replicates. * *p* = 0.025 S222A vs MEF2C, t-test. (E) Kaplan–Meier survival curves of primary recipient mice transplanted with wild-type *MLL-AF9; MEF2C*, dominant negative *MLL-AF9; MEF2C S222A* or control *MLL-AF9; MIT* transformed leukemias. N = 10 animals in each group. Kaplan–Meier survival curves of secondary recipient mice transplanted with (F) 10,000 cells, (G) 1,000 cells and (H) 10 cells from that shown in (E). N = 20 animals in each group. *p* = 0.022 S222A vs MEF2C for 10,000 cells; 4.34 × 10^−4^ S222A vs MEF2C for 1,000 cells; 4.4 × 10^−2^ S222A vs MEF2C for 10 cells, log rank test. qRT-PCR of (I) *Hdac7,* (J) *Nr4a1* and (K) *Jun* in primary *MLL-AF9; MEF2C* transformed cells. Error bars represent standard deviation of the mean for 3biological replicates.

**Figure S6.**
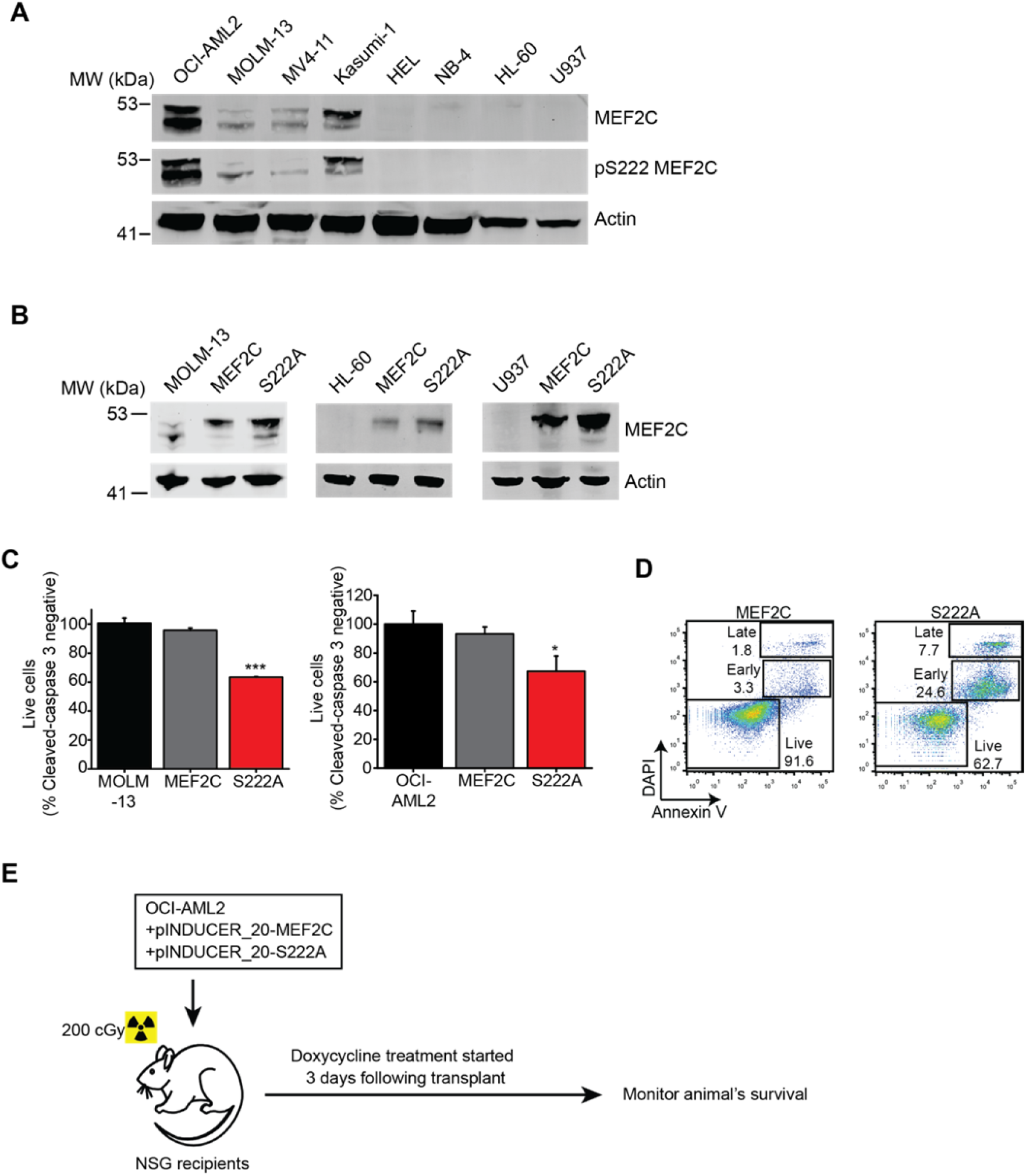
Related to Figure 3. MEF2C phosphorylation is required for leukemia maintenance and cell survival in MLL-r leukemia. (A) Western immunoblot analysis for MEF2C and pS222 MEF2C in human AML cell lines. (B) Western immunoblot analysis of MEF2C and pS222 MEF2C in MOLM-13, HL-60 and U937 cells lentivirally transduced with wild-type *MEF2C* or dominant negative *MEF2C S222A* transgenes and treated for 48 hours with 600 ng/ml doxycycline to induce transgene expression. (C) Cleaved–caspase 3 analysis in MOLM-13 and OCI-AML2 cells treated with 600 ng/ml doxycycline for 48 hours. Error bars represent standard deviation of the mean for 3 biological replicates. *** and * *p* = 7.8 × 10^-5^ and 1.3 × 10^-2^ S222A vs MEF2C for MOLM-13 and OCI-AML2, respectively (t-test). (D) Representative FACS plots of annexin V and DAPI staining in MOLM-13 cells treated with 600 ng/ml doxycycline for 48 hours. (E) Schematic for OCI-AML2 cells transplanted into NSG animals.

**Figure S7.**
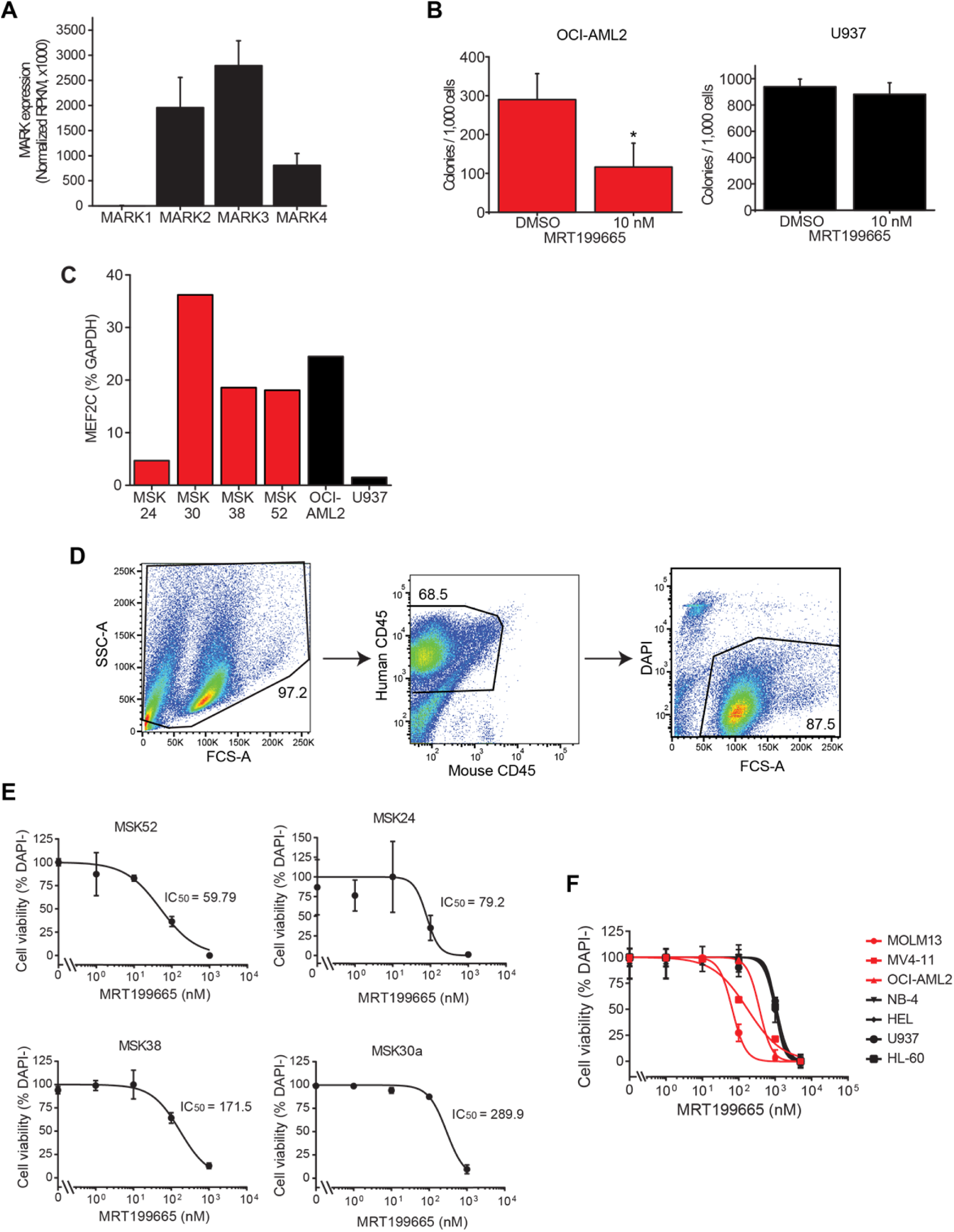
Related to Figure 5. Chemical inhibition of MARK kianses exhibits selective toxicity against MEF2C-activated human AML cells. (A) Gene expression of *MARK* isoforms in primary AML patient samples from The Cancer Genome Atlas (TCGA). Error bars represent standard deviation of the mean. (B) Colony formation of OCI-AML2 cells (red) and U937 cells (black) treated with 10 nM MRT199665 or DMSO vehicle control for 10 days. * *p*= 2.9 × 10^-2^ (t-test). Error bars represent standard deviation of the mean for 3 biological replicates. (C) qRT-PCR of *MEF2C* in primary AML specimens (red) compared to human AML cell lines (black). (D) Schematic of FACS analysis of patient derived xenograft (PDX) cells treated with MRT199665 for 48 hours. (E) Cell viability curves as a function of MRT199665 concentration of PDX samples treated for 48 hours with MRT199665. Error bars represent standard deviation of the mean for 3 biological replicates. (F) Cell viability curves as a function of MRT199665 concentration in human AML cell lines treated for 48 hours and assessed by FACS of DAPI viability staining in triplicate reactions. Error bars represent standard deviation of the mean for 3 biological replicates.

## SUPPLEMENTAL TABLES

**Table S1.** Demographic features and somatic mutations of induction failure and induction remission cohorts used for western blotting

| Patient ID | Therapy outcome | Gender | Age (Years) | Therapy | Somatic mutations | Genomic rearrangements |
| --- | --- | --- | --- | --- | --- | --- |
| MSK103 | R | F | 7.9 | ADE | NRAS_G12D; RUNX1_Q390fs*;<br>FLT3_R595_E596ins; | KMT2A duplication |
| MSK109 | R | F | 11.4 | ADE | WT1_S381fs*4; STK11_F354L;<br>FLT3_G5853_S583ins |  |
| MSK113 | R | M | 16.8 | ADE | ND |  |
| MSK110 | R | F | 13.8 | ADE | FLT3_T582_G583ins |  |
| MSK106 | R | M | 3.7 | ADE | KRAS_G12C; KIT_D816Y; NF2_splice<br>site 1739del | KMT2A_MLLT3 fusion |
| MSK148 | R | F | 17.1 | ADE | SETD2_R1625H; WT1_R369*;<br>MDM4_S367L; FLT3_E596_Y597ins | KMT2A duplication |
| MSK153 | R | F | 16.6 | ADE | U2AF1_R156H; YY1AP1_G412S | KMT2A_MLLT4 fusion |
| MSK156 | R | M | 13.0 | ADE | CEBPA_H24fs*; WT1_Q259*;<br>FLT3_V592-D593ins; EP300_splice;<br>CEBPA_T381_S319ins |  |
| MSK160 | R | M | 9.8 | ADE | FLT3_Q580-V581ins; CD36_N53fs*;<br>CEBPA_H24fs* |  |
| MSK161 | R | M | 6.6 | ADE | CEBPA_Q83fs*; WT1_E150*;<br>ZRSR2_R448-R449ins; CEBPA_T310-Q311ins |  |
| MSK30 | R | F | 67 | IDA+HDAC | FLT3_N676K, S584-S585ins;<br>DNMT3A_P244fs*; |  |
| MSK22 | R | M | 62 | 7+3 DNR<br>and AraC | NPM1_W288fs*; DNMT3A_R882H;<br>NRAS_G12D |  |
| MSK26 | R | F | 53 | 7+3 DNR<br>and AraC | DNMT3A_R882H; NPM1_W288fs*;<br>FLT3_Y599ins; TET2_C1281fs* |  |
| MSK02 | R | F | 44 | 7+3 DNR<br>and AraC | DNMT3A_R882H; EPHA5;R417Q;<br>NPM1_W288fs*; FLT3_G583-S584ins,<br>F590-Y591ins |  |
| MSK09 | R | F | 22 | 7+3 DNR<br>and AraC | NPM1_W288fs*; EPHA5_R919*;<br>TET2_H434fs*, W564*; FLT3_Y597- |  |
|  |  |  |  |  | E598ins |  |
| MSK104 | F | F | 13.5 | ADE | KRAS_G12C |  |
| MSK105 | F | M | 17.6 | ADE | FLT3_V579-Q580ins; WT1_A382fs* |  |
| MSK107 | F | M | 17.2 | ADE | DNMT3A_R882H; IDH2_R172K |  |
| MSK108 | F | F | 7.8 | ADE | FLT3_S574-Q575ins |  |
| MSK159 | F | M | 10.7 | ADE | PTPN11_D61Y; ASXL1_G645fs* |  |
| MSK130 | F | M | 0.8 | ADE | NRAS; Q61L; FLT2_D835H;<br>ICK_splice site del | KMT2A_MLLT10 fusion |
| MSK120 | F | M | 15.8 | ADE | FLT3_L610-E611ins; CD36_I399fs* |  |
| MSK138 | F | M | 6.8 | ADE | FLT3_T582-G583ins | NSD1_NUP98 fusion |
| MSK135 | F | M | 18.1 | ADE | FLT3_Y591-V592ins |  |
| MSK19 | F | M | 63 | 7+3 DNR<br>and AraC | DNMT3A_R882H; NPM1_W288fs*;<br>ARID2_Q1149fs*; FLT3_V592-D593ins |  |
| MSK27 | F | M | 55 | 7+3 DNR<br>and AraC | NPM1_W288fs*; IDH2_R140Q;<br>DNMT3A_splice site |  |
| MSK25 | F | M | 53 | 7+3 DNR<br>and AraC | DNMT3A_R882H; NPM1_W288fs*;<br>FLT3_R595-F596ins |  |
| MSK16 | F | F | 61 | 7+3 DNR<br>and AraC | GATA2_R362Q; NPM1_W288fs*;<br>FLT3_Y591ins |  |
| MSK04 | F | F | 50 | 7+3 DNR<br>and AraC | NPM1_W288fs*; FLT3_R595-E596ins |  |

Abbreviations: R, remission, F, failure; F, female, M, male; ADE, cytarabine, daunomycin and etoposide-based regimen; IDA, idarubicin; HDAC, high-dose cytarabine; DNR, daunorubicin.

**Table S2.** Antibodies

| Name | Clone | Catalog # | Source | Technique |
| --- | --- | --- | --- | --- |
| Rabbit anti-MEF2C | D80C1 | 5030 | Cell Signaling Technology (CST),<br>Danvers, MA, USA | WB |
| Rabbit anti-pS222 MEF2C | RRID:AB_2572427 | p1208-222 | PhosphoSolutions, | WB |
| Mouse anti-β-Actin | 8H10D10 | 3700 | CST | WB |
| Rabbit anti-MARK3 | -- | 9311 | CST | WB |
| Goat Anti Mouse IRDye 800W | -- | 926-32210 | Li-Cor, Nebraska, USA | WB |
| Goat anti-Rabbit IgG IRDye 680RD | -- | 925-68071 | Li-Cor | WB |
| APC anti-mouse B220 | RA3-6B2 | 103212 | BioLegend, San Diego, CA, USA | FC |
| FITC anti-mouse Gr-1 | RB6-8C5 | 11-5931-82 | eBioscience, San Diego, CA, USA | FC |
| PE-Cy5 anti-mouse CD11b | M1/70 | 15-0112-83 | eBioscience | FC |
| Pacific blue anti-mouse CD4 | RM4-5 | 558107 | BD Biosciences, San Jose, CA, USA | FC |
| Pe-Cy7 anti-mouse CD8 | 53-6.7 | 25-0081-81 | eBioscience | FC |
| Pacific Blue anti-mouse Sca-1 | D7 | 108120 | BioLegend | FC |
| APC anti-mouse CD117 | ACK2 | 17-1172-83 | eBioscience | FC |
| FITC anti-mouse CD34 | RAM34 | 11-0341-82 | eBioscience | FC |
| PE anti-mouse CD16/32 | 93 | 101308 | BioLegend | FC |
| FITC anti-mouse CD45.1 | A20 | 110705 | BioLegend | FC |
| Pe/Cy7 anti-mouse CD45.2 | 104 | 109829 | BioLegend | FC |
| FITC anti-mouse CD45 | 30-F11 | 103108 | BioLegend | FC |
| APC anti-human CD45 | HI30 | 304012 | BioLegend | FC |
| DAPI | -- | D1306 | Thermo Fisher Scientific, Waltham, MA | FC |
| APC Annexin V | -- | 640920 | BioLegend | FC |
| Alexa Fluor 647 Cleaved Caspase 3 | C92-605 | 560626 | BD Biosciences | FC |

Abbreviations: WB, western blot; FC, flow cytometry.

## INDEX FOR SUPPLEMENTAL DATA

Supplemental data files assembled in Brown_etal_SupplementaryData.zip

| File number | File name | Description |
| --- | --- | --- |
| 1 | Phosphoproteomics_ITRAQ_Summary.xlsx | Table of phosphopeptides identified in a screen for primary induction failure compared to complete remission AML |
| 2 | RNAseq_DEG_S222A+_S222A-.xlsx | List of differentially expressed genes between S222A+ and S222A- cells by RNA sequencing |
| 3 | OCIAML2_GSEA_report_down.xlsx | Table of gene sets downregulated in S222A MEF2C compared to MEF2C in OCIAML2 cells by GSEA |
| 4 | OCIAML2_GSEA_report_up.xlsx | Table of gene sets upregulated in S222A MEF2C compared to MEF2C in OCIAML2 cells by GSEA |
| 5 | Serine threonine kinase screen report.xls | Table of results for recombinant protein kinase screen for pS222 MEF2C |

## SUPPLEMENTAL EXPERIMENTAL PROCEDURES

### Reagents

All reagents were obtained from Sigma Aldrich unless otherwise stated. Synthetic oligonucleotides were obtained from Eurofins MWG Operon (Hunstville, AL, USA) and purified by HPLC. Synthetic ultramer oligonucleotides for CRISPR/Cas9 mouse genome engineering were obtained from Integrated DNA Technologies (Coralville, IA). DNA Sanger sequencing was performed by Genewiz (South Plainfield, NJ, USA).

### Generation of *Mef2c*^*S222A*^ and *Mef2c*^*S222D*^ Mice

All mouse studies were conducted with approval from the Memorial Sloan Kettering Cancer Center Institutional Animal Care and Use Committee. C57BL/6J, NOD-Scid IL2Rγ-null (NSG) and B6. SJL-Ptprca Pepcb/BoyJ mice were all obtained from Jackson Labs (Bar Harbor, ME, USA). Generation of knock-in *Mef2c*^*S222A*^ and *Mef2c*^*S222D*^ mutant mice using CRISPR/Cas9 genome editing and homologous recombination is described in the Supplemental Experimental Procedures. (Wang et al., 2013). A CRISPR gRNA targeting exon 5 of the *Mef2c* gene was designed (GGAATGGATACGGCAACCCC) and expressed *in vitro* using the approach previously described (Romanienko et al., 2016). The donor templates for recombination were designed according to Wefers and colleagues (Wefers et al., 2014). For *Mef2c*^*S222A*^, we used the oligonucleotide

ATATGAGCTGTTCTTTAAAAACATGCACACGCCTCGATTTCTCCCTGAGTTTTGTCATCTCT TGTGTTAACAGGGAATGGATACGGCAACCC<u>CCG**c**</u>AAC<u>**g**C**g**</u>CCAGGCCTGCTGGTCTCACCTGGTAACCTGAACAAGAATATACAAGCCAAATCTCCTCCCCCTATGAATCTAGGAATGAATA ATCGTA, where lowercase nucleotides indicate in-frame substitutions from the reference sequence and underlined regions indicates the addition of a HhaI and AciI restriction sites, respectively. For *Mef2c*^*S222D*^, we used the oligonucleotide ATATGAGCTGTTCTTTAAAAACATGCACACGCCTCGATTTCTCCCTGAGTTTTGTCATCTCTTGTGTTAACAGGGAATGGATACGGCAACCC<u>CCG**c**</u>AAC<u>**gat**C</u>CAGGCCTGCTGGTCTCACCTGGTAACCTGAACAAGAATATACAAGCCAAATCTCCTCCCCCTATGAATCTAGGAATGAATAA TCGTA, where lowercase nucleotides indicate in-frame substitutions from the reference sequence and underlined regions indicates the addition of a Sau3A and AciI restriction sites, respectively. Mouse zygotes from C57BL/6J:CBA F1 hybrids were injected with 100 ng/μl of gRNA, 50 ng/μl Cas9 mRNA and 50 ng/μl of donor oligonucleotide into the pronucleus. Founder mice were first screened using restriction endonuclease digestion of genomic DNA isolated from tail tissues. Digestions were carried out using HhaI and AciI for Mef2cS222A and Sau3A and AciI for Mef2cS222D alleles. The positive mice were then analyzed by DNA sequencing of PCR products obtained from amplifying the targeted region of exon 5 using the following forward and reverse primers, respectively: GTATGACTCACAGCTAGGCGTATC, and CATCGTGTTCTTGCTGCCAGGTG. For all experiments using *Mef2c* knock-in mutant animals, wild-type litter mates were used as controls.

### Plasmids and expression vectors

Human full-length *MEF2C* cDNA isoform 1 (RefSeq ID: NM_002397.4) was cloned into the pMSCV-IRES-tdTomato (MIT) retroviral expression vector, kindly obtained from Scott Armstrong, (Zhu et al., 2016), to generate *MIT*-*MEF2C*. pRecLV105 lentiviral vector (GeneCopoeia, Rockville, MD, USA) was used to generate pRecLV105-*MEF2C*. The doxycycline-inducible pINDUCER20 lentiviral expression vector was a kind gift from Thomas Westbrook (Meerbrey et al., 2011), and used to generate pINDUCER20-*MEF2C* using Gateway cloning according to the manufacturer’s instructions (Fisher Scientific, Watham, MA, USA). Site-directed mutagenesis was used to generate *MEF2C S222A* according to the manufacturer’s instructions (QuikChange Lightning, Agilent, Santa Clara, CA, USA). Human full-length *MARK3* cDNA isoform c (RefSeq ID: NM_2376.6) was amplified from the OCI-AML2 human AML cell line and cloned using Gateway cloning into pLX304. (Yang et al., 2011). Lentivirus packaging vectors psPAX2 (Addgene plasmid #12260) and pMD2.G (Addgene plasmid #12259) were a gift from Didier Trono, and ecotropic retroviral packaging vector pCL-Eco was a gift from Inder Verma (Addgene plasmid #12371) (Naviaux et al., 1996). The MSCV-IRES-GFP *MLL-AF9* expression vector was a kind obtained from Scott Armstrong (Krivtsov et al., 2006).

### Cell culture

Human AML cell lines were obtained from DSMZ (Brunswick, Germany) and cultured in RPMI-1640 medium supplemented with 10% fetal bovine serum (FBS) and antibiotics (100 U / ml penicillin and 100 μg / ml streptomycin) in a humidified atmosphere at 37 °C and 5% CO_2_. HEK293T cells were obtained from American Type Culture Collection (ATCC, Manassas, Virginia, USA) and cultured in Dulbecco’s Modified Eagle medium (DMEM) supplemented with 10% FBS and antibiotics (100 U / ml penicillin and 100 μg / ml streptomycin). All cell lines were authenticated by STR genotyping (Genetica DNA Laboratories, Burlington, NC, USA) and the absence of *Mycoplasma sp.* contamination was verified using Lonza MycoAlert (Lonza Walkersville, Inc., Walkersville, MD, USA). Short-term culture of patient AML specimens were performed by thawing viably preserved cells in IMDM supplemented with 10% FBS, 0.1 mM β-mercaptoethanol, and 100 ng/ml human SCF, 20 ng/ml human IL3, 20 ng/ml human G-CSF, and 50 ng/ml human FLT3-L (all obtained from Peprotech, Rocky Hill, New Jersey, USA).

### Virus production and cell transduction

Lentiviral production was performed as previously described (Kentsis et al., 2012). Briefly, HEK293T packaging cells were transfected with TransIT-LTI according to the manufacturer’s instructions (Mirus Bio LLC, Madison, WI, USA). For lentiviral expression vectors, a 2:1:1 ratio of the expression vector, packaging and pseudotyping plasmids was used. For retroviral expression vectors, a 1:1 ratio of expression vector to pCL-Eco packaging plasmidwas used. Viral supernatant was collected at 48 and 72 hours post-transfection, pooled, filtered, concentrated by centrifugation using the Amicon Ultra-15 Centrifugal Filter Units (EMD Millipore, Darmstadt, Germany) and stored at -80 °C. HEK293T cells were transduced by spin infection with pRecLV105 or pLX304 viral particles at a multiplicity of infection (MOI) of 1 in the presence of 8 µg/ml polybrene, and transduced cells were selected with 2 µg/ml puromycin for 3 days (for pRecLV105) or 10 µg/ml blasticidin (for pLX304) for 8 days. Human AML cell lines were transduced by spin infection with pINDUCER20 viral particles at a MOI of 1, and transduced cells were selected with 2 mg/ml G418 sulfate for 14 days. Mouse hematopoietic progenitor cells were transduced with viral particles at MOI of 20 by spin infection on retronectin-treated tissue culture plates (Takara Bio USA, Inc; Mountain View, CA, USA) and GFP or tdTomato positive cells were isolated by FACS 3 days after transduction.

### Isolation of mouse hematopoietic progenitor cells

Femurs and tibias were dissected from 8-12 week-old C57BL/6J or transgenic mice, and cells were extracted using mortar and pestle crushing, and mesh filtration. Red blood cells were lysed using the RBC Lysis Buffer according to the manufacturer’s instructions (BioLegend, San Diego, CA, USA). Progenitor cells were isolated using lineage depletion with the Mouse Lineage Depletion Kit according to the manufacturer’s instructions (MACS Miltenyi Biotech, Bergisch Gladbach, Germany). Isolated cells were subsequently stained with the following antibodies for isolation of granulocyte/macrophage progenitor cells (GMP): Sca-1, CD16/32, CD117 and CD34. Fluorescence-activated cell sorting was performed using the BD FACSAria (Becton Dickinson, San Jose, CA) and FACS analysis was done using FlowJo software (FlowJo, Ashland, Oregon, USA). Sorted GMP cells were cultured overnight in IMDM supplemented with 15% FBS, 100 U / ml penicillin and 100 μg / ml streptomycin, and 50 ng/ml mouse SCF, 10 ng/ml mouse IL-3, and 10 ng/ml mouse IL-6 (all from Peprotech). Antibodies used in flow cytometry are detailed in Table S2.

### Leukemia transplants

For primary *MLL-AF9* mouse leukemia transplants, 150,000 transduced cells were transplanted by intravenous injection into lethally irradiated (900 rad) C57BL/6J animals with 125,000 support bone marrow cells and monitored for the development of leukemia. For secondary transplants, leukemic cells harvested from the bone marrow of moribund animals were transplanted by intravenous injection into sub-lethally irradiated (600 rad) C57BL/6J mice at cell doses indicated throughout the text. Leukemia stem cell frequency was determined using ELDA software (Hu and Smyth, 2009).

For competitive bone marrow transplantation studies, 1 million CD45.2 mononuclear bone marrow cells from *MEF2C* transgenic mice were mixed with 1 million CD45.1 mononuclear bone marrow cells from B6.SJL-Ptprca Pepcb/BoyJ mice and transplanted via tail vein injection in lethally irradiated C57BL/6J recipient animals. Transplanted mice were monitored by CD45.1/CD45.2 peripheral blood chimerism using FACS with the surface markers CD45.2 and CD45.1.

For human xenograft transplants, NSG mice (8-10 weeks old) were sub-lethally irradiated (200 rad) and transplanted with 500,000 OCI-AML2 cells via tail vein injection. Doxycycline-inducible transgene expression was induced using doxycycline chow 3 days post-transplantation (Harlan, Indianapolis, USA).

### Cell proliferation studies

For human AML cell lines transduced with pINDUCER20-*MEF2C* and pINDUCER-*MEF2C-S222A*, 1 × 10^4^ cells were incubated in 100 μl with 600 ng/ml of doxycycline for 72 hours in 96-well plates. For chemotherapy treatments, cytarabine and doxorubicin dissolved in PBS were added to cells after 24 hours of doxycycline treatment. Cell growth was assessed using the Cell Titer-Glo Luminescent Viability assay (Promega, Madison, WI, USA), as measured using luminescence on the Infinite M1000Pro plate reader using integration time of 250 milliseconds (Tecan, Männedorf, Switzerland). For assays using patient AML cells, cells were assayed by FACS using the BD FACSCanto II cytometer (BD Biosciences) after staining cells with human CD45, mouse CD45 and DAPI viability stain. Antibodies used in flow cytometry are detailed in Table S2.

### Clonogenic assays

All colony forming assays were performed using HSC001 methylcellulose medium, according to the manufacturer’s instructions (R&D Systems, Minneapolis, MN, USA). For GMP cells transduced with *MLL-AF9*, 1,000 cells/replicate were plated in 1.27% final concentration methylcellulose with IMDM supplemented with 10% FBS and mouse cytokines (10 ng/ml IL-3, 10ng/ml IL-6 and 20 ng/ml SCF). Leukemia blast colonies were counted after 7 days. For re-plating experiments, cells were pooled, counted and replated at the same density. Purified GMP cells (20,000 cells/replicate) from the *Mef2c*^*S222A*^ and *Mef2c*^*S222D*^ transgenic mice were plated in 1.27% final concentration methylcellulose with IMDM supplemented with 10% FBS and mouse cytokines (10 ng/ml IL-3, 10 ng/ml IL-6 and 50 ng/ml SCF). Colonies from CFU-G, CFU-G and CFU-GM units were counted after 14 days.

### Transcriptional reporter assays

MEF2C transcriptional activity was quantified using the Cignal MEF2 reporter assay, using the CTAGCGCTCTAAAAATAACCCT response element, according to the manufacturer’s instructions (Qiagen, Hilden, Germany). Cells were transfected with the MEF2C reporter gene using the Fugene transfection reagent (Promega) and detected using the Dual-Glo Luciferase Assay System (Promega), according to the manufacturer’s instructions. MEF2C transcriptional activity was calculated using the ratios of the firefly/*Renilla* signals, and normalized to the internal positive control for the reporter assay.

### Recombinant kinase screen

The protein kinase screen was performed using a recombinant serine kinase library as previously described (Kitagawa et al., 2012). Briefly, 172 serine/threonine kinases (Table S4) were individually expressed as N-terminal GST-fusion proteins in insect cells, and purified using glutathione sepharose chromatography. A synthetic peptide corresponding to phosphoserine 222 for human MEF2C (RefSeq ID: NM_002397.4) Ac-GNPRN[pS]PGLLVC-NH2 was synthesized (Tufts University, Medford, MA, USA), purified by HPLC, and confirmed by mass spectrometry. For the screen, the MEF2C peptide was dissolved in DMSO at 10 μM, 1 μM, and 0.1 μM, and specific kinase activity on respective substrates was determined by the off-chip mobility shift assay (MSA) using the LabChipTM3000 instrument (Caliper Life Sciences, Hopkinton, MA, USA). The kinase reaction was analyzed by the product ratio, which was calculated from the peak heights of the kinase substrate (K) and MEF2C (M) peptides (K/(K+M)). Additional protein kinases were assayed using the immobilized metal ion affinity-based fluorescence polarization (IMAP) system, where the kinase reaction was evaluated by the fluorescence polarization at 485 nm for excitation and 530 nm for emission. Staurosporine was used as the positive control for almost all kinases and the readout of this was set as 0% inhibition, with the readout value of no kinase set as a 100% inhibition. Inhibition of 20% or more was used to score positive signals.

### Western immunoblotting

Primary human or mouse leukemia cells were lysed in 100 μl of lysis buffer containing 30 mM Tris HCl, 1% sodium dodecyl sulfate (SDS), 7% glycerol, 1.25% beta-mercaptoethanol, 0.2 mg/ml Bromophenol Blue, pH 6.8 per 1 million cells and incubated at 95 °C for 10 minutes. Human AML cell lines and HEK293T cells were lysed in RIPA buffer (15 mM NaCl, 1% NP-40, 0.5% sodium deoxycholate, 0.1% SDS, 50 mM Tris, pH 8.0) and sonicated using the Covaris S220 adaptive focused sonicator, according to the manufacturer’s instructions (Covaris, Woburn, CA). Lysates were cleared by centrifugation at 11,000 × g for 10 minutes at 4 °C and clarified lysates were denatured in Laemmli sample buffer at 95 °C for 10 minutes. For phosphatase treatment, cell lines were lysed in RIPA buffer without SDS and treated with 800U λ phosphatase (P9614) and 6500U alkaline phosphatase (P0114) for 30 minutes before the addition of 1% SDS. Lysates were separated by sodium dodecyl sulfate polyacrylamide gel electrophoresis (SDS-PAGE) and transferred onto an Immobilon-FLPVDF membrane (EMD Millipore, Darmstadt, Germany). Membranes were blocked with the Odyssey Blocking Buffer (Li-Cor, Nebraska, USA) and blotted with primary antibodies for MEF2C (1:1000), pS222 MEF2C (1:1000), MARK3 (1:1000) or beta-actin (1:1000), followed by blotting with goat IRDye 680 RD anti-rabbit and IRDye 800 CW anti-mouse secondary antibodies. For peptide competition assays, pS222 MEF2C was pre-blocked with synthetic peptides for phospho-MEF2C and nonphospho-MEF2C corresponding to amino acids surrounding S222. Signals were was recorded and quantified using the Odyssey CLx fluorescence scanner (Li-Cor, Nebraska, USA). Antibodies used in western blot are detailed in Table S2.

### Flow cytometry

Antibodies used in flow cytometry are detailed in Table S2. For analysis of annexin V, cells were washed and resuspended in annexin V binding buffer (10 mM HEPES, pH 7.4, 140 NaCl, 2.5 mM CaCl_2_). Cells were stained with annexin V for 15 minutes at room temperature in the dark, with the addition of DAPI before analysis by flow cytometry. For analysis of caspase 3 cleavage, cells were fixed with BD Cytofix/Cytoperm buffer according to the manufacturer’s protocol (BD Biosciences). Fixed cells were stained with Alexa Fluor 647-conjugated anti cleaved caspase-3 (BD Biosciences) in BD Perm/Wash Buffer (BD Biosciences) for 30 minutes at room temperature in the dark. Bone marrow mononuclear cells were assessed using the surface cell marker antibodies B220, Gr-1, CD11b, CD4 and CD8. Flow cytometry was analyzed on a BD LSRFortessa and FlowJo software.

### Giemsa staining

Dip Quick Stain (J-322, Jorgensen Laboratories, Inc) was used according to the manufacturer’s protocol for Giemsa staining of peripheral blood smears and bone marrow cytospins. Cytospins of bone marrow cells were prepared using a benchtop Cytospin Centrifuge (Thermo Fisher Scientific) of 2 × 10^5^ cells in PBS onto Superfrost Plus microscope slides, according to the manufacturer’s instructions (Thermo Fisher Scientific).

### Quantitative RT-PCR

RNA was extracted using TRIzol Reagent (Thermo Fisher Scientific, Waltham, MA) followed by the Qiagen RNeasy Mini Kit according to the manufacturer’s instructions. Complementary DNA was synthesized using the SuperScript III First-Strand Synthesis system according to the manufacturer’s instructions (Invitrogen). Quantitative real-time PCR was performed using the KAPA SYBR FAST PCR polymerase, according to the manufacturer’s instructions (Kapa Biosystems, Wilmington, MA, USA). Ct values were calculated using ROX normalization on the ViiA 7 software (Applied Biosystems).

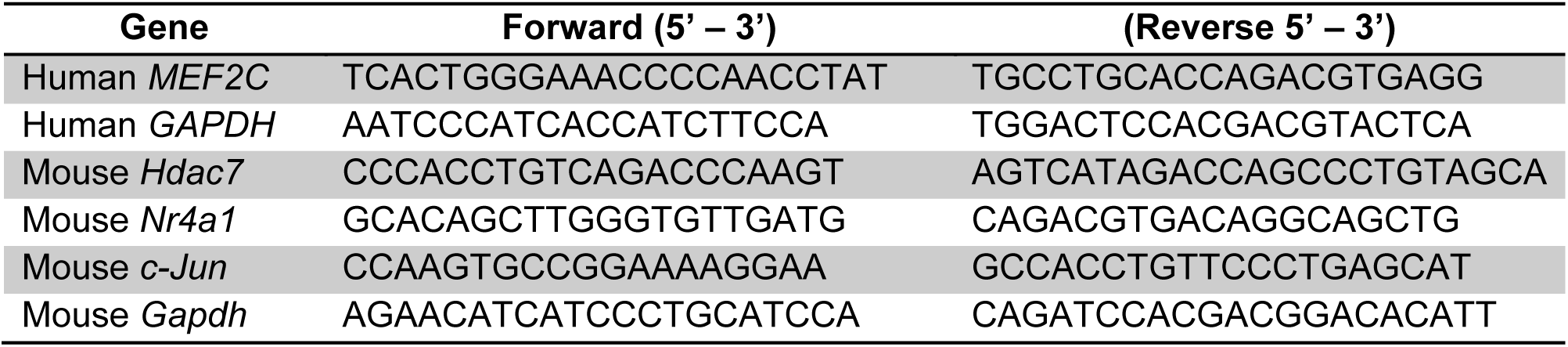
Sequences for RT-qPCR primers

### Gene expression profiling

RNA was extracted from cells as described above. The quality of RNA was verified on the Agilent Bioanalyzer 2100 platform (Agilent, Santa Clara, CA, USA). For RNA sequencing, a Poly-A tail library selection was performed. Sequencing was done using the Illumina Next-Gen Sequencing HiSeq platform (Illumina, San Diego, CA) with 30-45 million 50bp, paired-end reads.

### Data Analysis

RNA-seq raw reads were quality and adapter trimmed using ‘trim_galore’ before aligning to human assembly hg19 with STAR v2.5 using the default parameters. Coverage and post-alignment quality were assessed using the Picard tool CollectRNASeqMetrics (http://broadinstitute.github.io/picard/). Read count tables were created using HTSeq v0.6.1. Normalization and expression dynamics were evaluated with DESeq2 using the default parameters. Gene set enrichment analysis (GSEA, http://software.broadinstitute.org/gsea/) was performed with the pre-ranked option and default parameters. The log2 fold change from DESeq2 was used as input, and tested against all gene sets in MSigDB version 5.1. For *MARK1-4* gene expression, raw sequences of transcriptome data collected as part of the Cancer Genome Atlas (TCGA) from 138 patients with AML (Cancer Genome Atlas Research, 2013) were aligned to the NCBI37/mm9 human reference genome using STAR and differentialexpression analysis obtained using the DESeq2 Bioconductor algorithm. Proteomics and genomics data have been deposited to MassIVE (https://massive.ucsd.edu/) and Gene Expression Omnibus (https://www.ncbi.nlm.nih.gov/geo/), respectively.

## REFERENCES

Barneda-Zahonero, B., Roman-Gonzalez, L., Collazo, O., Rafati, H., Islam, A. B., Bussmann, L. H., di Tullio, A., De Andres, L., Graf, T., Lopez-Bigas, N., et al. (2013). HDAC7 is a repressor of myeloid genes whose downregulation is required for transdifferentiation of pre-B cells into macrophages. PLoS genetics 9, e1003503.

Breems, D. A., Van Putten, W. L., Huijgens, P. C., Ossenkoppele, G. J., Verhoef, G. E., Verdonck, L. F., Vellenga, E., De Greef, G. E., Jacky, E., Van der LelieJ., et al. (2005). Prognostic index for adult patients with acute myeloid leukemia in first relapse. Journal of clinical oncology: official journal of the American Society of Clinical Oncology 23, 1969–1978.

Brown, F. C., Cifani, P., Drill, E., He, J., Still, E., Zhong, S., Balasubramanian, S., Pavlick, D., Yilmazel, B., Knapp, K. M., et al. (2017). Genomics of primary chemoresistance and remission induction failure in paediatric and adult acute myeloid leukaemia. British journal of haematology 176, 86–91.

Burnett, A., Wetzler, M., and Lowenberg, B. (2011). Therapeutic advances in acute myeloid leukemia. Journal of clinical oncology: official journal of the American Society of Clinical Oncology 29, 487–494.

Cancer Genome Atlas Research, N (2013) Genomic and epigenomic landscapes of adult de novo acute myeloid leukemia. The New England journal of medicine 368, 2059–2074.

Choe, L., D’Ascenzo, M., Relkin, N. R., Pappin, D., Ross, P., Williamson, B., Guertin, S., Pribil, P., and Lee, K. H. (2007). 8-plex quantitation of changes in cerebrospinal fluid protein expression in subjects undergoing intravenous immunoglobulin treatment for Alzheimer’s disease. Proteomics 7, 3651–3660.

Clark, K., MacKenzie, K. F., Petkevicius, K., Kristariyanto, Y., Zhang, J., Choi, H. G., Peggie, M., Plater, L., Pedrioli, P. G., McIver, E., et al. (2012). Phosphorylation of CRTC3 by the salt-inducible kinases controls the interconversion of classically activated and regulatory macrophages. Proceedings of the National Academy of Sciences of the United States of America 109, 16986–16991.

Coombs, C. C., Tallman, M. S., and Levine, R. L. (2016). Molecular therapy for acute myeloid leukaemia. Nat Rev Clin Oncol 13, 305–318.

Rooij, de J. D.Zwaan, C. M.Heuvel-Eibrink, van den M. (2015). Pediatric AML: From Biology to Clinical Management. Journal of clinical medicine 4, 127–149.

Ding, L., Ley, T. J., Larson, D. E., Miller, C. A., Koboldt, D. C., Welch, J. S., Ritchey, J. K., Young, M. A., Lamprecht, T., McLellan, M. D., et al. (2012). Clonal evolution in relapsed acute myeloid leukaemia revealed by whole-genome sequencing. Nature 481, 506–510.

Du, Y., Spence, S. E., Jenkins, N. A., and Copeland, N. G. (2005). Cooperating cancer-gene identification through oncogenic-retrovirus-induced insertional mutagenesis. Blood 106, 2498–2505.

Farrar, J. E., Schuback, H. L., Ries, R. E., Wai, D., Hampton, O. A., Trevino, L. R., Alonzo, T. A., Guidry Auvil, J. M., Davidsen, T. M., Gesuwan, P., et al. (2016). Genomic Profiling of Pediatric Acute Myeloid Leukemia Reveals a Changing Mutational Landscape from Disease Diagnosis to Relapse. Cancer research 76, 2197–2205.

Ficarro, S. B., Adelmant, G., Tomar, M. N., Zhang, Y., Cheng, V. J., and Marto, J. A. (2009). Magnetic bead processor for rapid evaluation and optimization of parameters for phosphopeptide enrichment. Analytical chemistry 81, 4566–4575.

Ficarro, S. B., Biagi, J. M., Wang, J., Scotcher, J., Koleva, R. I., Card, J. D., Adelmant, G., He, H., Askenazi, M., Marshall, A. G., et al. (2014). Protected amine labels: a versatile molecular scaffold for multiplexed nominal mass and sub-Da isotopologue quantitative proteomic reagents. Journal of the American Society for Mass Spectrometry 25, 636–650.

Ficarro, S. B., Zhang, Y., Carrasco-Alfonso, M. J., Garg, B., Adelmant, G., Webber, J. T., Luckey, C. J., and Marto, J. A. (2011). Online nanoflow multidimensional fractionation for high efficiency phosphopeptide analysis. Molecular & cellular proteomics: MCP 10, 111 011064.

Goardon, N., Marchi, E., Atzberger, A., Quek, L., Schuh, A., Soneji, S., Woll, P., Mead, A., Alford, K. A., Rout, R., et al. (2011). Coexistence of LMPP-like and GMP-like leukemia stem cells in acute myeloid leukemia. Cancer cell 19, 138–152.

Gu, Z., Churchman, M., Roberts, K., Li, Y., Liu, Y., Harvey, R. C., McCastlain, K., Reshmi, S. C., Payne-Turner, D., Iacobucci, I., et al. (2016). Genomic analyses identify recurrent MEF2D fusions in acute lymphoblastic leukaemia. Nature communications 7, 13331.

Guo, S., and Kemphues, K. J. (1995). par-1, a gene required for establishing polarity in C. elegans embryos, encodes a putative Ser/Thr kinase that is asymmetrically distributed. Cell 81, 611–620.

Guryanova, O. A., Shank, K., Spitzer, B., Luciani, L., Koche, R. P., Garrett-Bakelman, F. E., Ganzel, C., Durham, B. H., Mohanty, A., Hoermann, G., et al. (2016). DNMT3A mutations promote anthracycline resistance in acute myeloid leukemia via impaired nucleosome remodeling. Nature medicine 22, 1488–1495.

Herglotz, J., Unrau, L., Hauschildt, F., Fischer, M., Kriebitzsch, N., Alawi, M., Indenbirken, D., Spohn, M., Muller, U., Ziegler, M., et al. (2016). Essential control of early B-cell development by Mef2 transcription factors. Blood 127, 572–581.

Hollink, I. H., van den Heuvel-Eibrink, M. M., Arentsen-Peters, S. T., Pratcorona, M., Abbas, S., Kuipers, J. E., Van Galen, J. F., Beverloo, H. B., Sonneveld, E., Kaspers, G. J., et al. (2011). NUP98/NSD1 characterizes a novel poor prognostic group in acute myeloid leukemia with a distinct HOX gene expression pattern. Blood 118, 3645–3656.

Hope, K. J., Jin, L., and Dick, J. E. (2004). Acute myeloid leukemia originates from a hierarchy of leukemic stem cell classes that differ in self-renewal capacity. Nature immunology 5, 738–743.

Hurov, J. B., Huang, M., White, L. S., Lennerz, J., Choi, C. S., Cho, Y. R., Kim, H. J., Prior, J. L., Piwnica-Worms, D., Cantley, L. C., et al. (2007). Loss of the Par-1b/MARK2 polarity kinase leads to increased metabolic rate, decreased adiposity, and insulin hypersensitivity in vivo. Proceedings of the National Academy of Sciences of the United States of America 104, 5680–5685.

Kang, J., Gocke, C. B., and Yu, H. (2006). Phosphorylation-facilitated sumoylation of MEF2C negatively regulates its transcriptional activity. BMC biochemistry 7, 5.

Kasimir-Bauer, S., Ottinger, H., Meusers, P., Beelen, D. W., Brittinger, G., Seeber, S., and Scheulen, M. E. (1998). In acute myeloid leukemia, coexpression of at least two proteins, including P-glycoprotein, the multidrug resistance-related protein, bcl-2, mutant p53, and heat-shock protein 27, is predictive of the response to induction chemotherapy. Experimental hematology 26, 1111–1117.

Kentsis, A., Reed, C., Rice, K. L., Sanda, T., Rodig, S. J., Tholouli, E., Christie, A., Valk, P. J., Delwel, R., Ngo, V., et al. (2012). Autocrine activation of the MET receptor tyrosine kinase in acute myeloid leukemia. Nature medicine 18, 1118–1122.

Kihara, R., Nagata, Y., Kiyoi, H., Kato, T., Yamamoto, E., Suzuki, K., Chen, F., Asou, N., Ohtake, S., Miyawaki, S., et al. (2014). Comprehensive analysis of genetic alterations and their prognostic impacts in adult acute myeloid leukemia patients. Leukemia 28, 1586–1595.

Kim, H., Lee, J. E., Kim, B. Y., Cho, E. J., Kim, S. T., and Youn, H. D. (2005). Menin represses JunD transcriptional activity in protein kinase C theta-mediated Nur77 expression. Exp Mol Med 37, 466–475.

Kitagawa, D., Gouda, M., Kirii, Y., Sugiyama, N., Ishihama, Y., Fujii, I., Narumi, Y., Akita, K., and Yokota, K. (2012). Characterization of kinase inhibitors using different phosphorylation states of colony stimulating factor-1 receptor tyrosine kinase. Journal of biochemistry 151, 47–55.

Klco, J. M., Miller, C. A., Griffith, M., Petti, A., Spencer, D. H., Ketkar-Kulkarni, S., Wartman, L. D., Christopher, M., Lamprecht, T. L., Helton, N. M., et al. (2015). Association Between Mutation Clearance After Induction Therapy and Outcomes in Acute Myeloid Leukemia. Jama 314, 811–822.

Kosuga, S., Tashiro, E., Kajioka, T., Ueki, M., Shimizu, Y., and Imoto, M. (2005). GSK-3beta directly phosphorylates and activates MARK2/PAR-1. The Journal of biological chemistry 280, 42715–42722.

Krivtsov, A. V., Twomey, D., Feng, Z., Stubbs, M. C., Wang, Y., Faber, J., Levine, J. E., Wang, J., Hahn, W. C., Gilliland, D. G., et al. (2006). Transformation from committed progenitor to leukaemia stem cell initiated by MLL-AF9. Nature 442, 818–822.

Laszlo, G. S., Alonzo, T. A., Gudgeon, C. J., Harrington, K. H., Kentsis, A., Gerbing, R. B., Wang, Y. C., Ries, R. E., Raimondi, S. C., Hirsch, B. A., et al. (2015). High expression of myocyte enhancer factor 2C (MEF2C) is associated with adverse-risk features and poor outcome in pediatric acute myeloid leukemia: a report from the Children’s Oncology Group. Journal of hematology & oncology 8, 115.

Li, H., Radford, J. C., Ragusa, M. J., Shea, K. L., McKercher, S. R., Zaremba, J. D., Soussou, W., Nie, Z., Kang, Y. J., Nakanishi, N., et al. (2008). Transcription factor MEF2C influences neural stem/progenitor cell differentiation and maturation in vivo. Proceedings of the National Academy of Sciences of the United States of America 105, 9397–9402.

Lin, Q., Schwarz, J., Bucana, C., and Olson, E. N. (1997). Control of mouse cardiac morphogenesis and myogenesis by transcription factor MEF2C. Science 276, 1404–1407.

Lu, J., McKinsey, T. A., Nicol, R. L., and Olson, E. N. (2000). Signal-dependent activation of the MEF2 transcription factor by dissociation from histone deacetylases. Proceedings of the National Academy of Sciences of the United States of America 97, 4070–4075.

Lu, Y., Loh, Y. H., Li, H., Cesana, M., Ficarro, S. B., Parikh, J. R., Salomonis, N., Toh, C. X., Andreadis, S. T., Luckey, C. J., et al. (2014). Alternative splicing of MBD2 supports self-renewal in human pluripotent stem cells. Cell stem cell 15, 92–101.

Ma, K., Chan, J. K., Zhu, G., and Wu, Z. (2005). Myocyte enhancer factor 2 acetylation by p300 enhances its DNA binding activity, transcriptional activity, and myogenic differentiation. Molecular and cellular biology 25, 3575–3582.

Mao, Z., Bonni, A., Xia, F., Nadal-Vicens, M., and Greenberg, M. E. (1999). Neuronal activity-dependent cell survival mediated by transcription factor MEF2. Science 286, 785–790.

McKinsey, T. A., Zhang, C. L., and Olson, E. N. (2002). MEF2: a calcium-dependent regulator ofcell division, differentiation and death. Trends in biochemical sciences 27, 40–47.

Miller, P. G., AL-Shahrour, F., Hartwell, K. A., Chu, L. P., Jaras, M., Puram, R. V., Puissant, A., Callahan, K. P., Ashton, J., McConkey, M. E., et al. (2013). In Vivo RNAi screening identifies a leukemia-specific dependence on integrin beta 3 signaling. Cancer cell 24, 45–58.

Molkentin, J. D., Black, B. L., Martin, J. F., and Olson, E. N. (1996). Mutational analysis of the DNA binding, dimerization, and transcriptional activation domains of MEF2C. Molecular and cellular biology 16, 2627–2636.

Nagel, S., Meyer, C., Quentmeier, H., Kaufmann, M., Drexler, H. G., and MacLeod, R. A. (2008). MEF2C is activated by multiple mechanisms in a subset of T-acute lymphoblastic leukemia cell lines. Leukemia 22, 600–607.

Nesic, D., Miller, M. C., Quinkert, Z. T., Stein, M., Chait, B. T., and Stebbins, C. E. (2010). Helicobacter pylori CagA inhibits PAR1-MARK family kinases by mimicking host substrates. Nature structural & molecular biology 17, 130–132.

Pan, R., Hogdal, L. J., Benito, J. M., Bucci, D., Han, L., Borthakur, G., Cortes, J., DeAngelo, D. J., Debose, L., Mu, H., et al. (2014). Selective BCL-2 inhibition by ABT-199 causes on-target cell death in acute myeloid leukemia. Cancer discovery 4, 362–375.

Papaemmanuil, E., Gerstung, M., Bullinger, L., Gaidzik, V. I., Paschka, P., Roberts, N. D., Potter, N. E., Heuser, M., Thol, F., Bolli, N., et al. (2016). Genomic Classification and Prognosis in Acute Myeloid Leukemia. The New England journal of medicine 374, 2209–2221.

Parikh, J. R., Askenazi, M., Ficarro, S. B., Cashorali, T., Webber, J. T., Blank, N. C., Zhang, Y., and Marto, J. A. (2009). multiplierz: an extensible API based desktop environment for proteomics data analysis. BMC Bioinformatics 10, 364.

Pierceall, W. E., Kornblau, S. M., Carlson, N. E., Huang, X., Blake, N., Lena, R., Elashoff, M., Konopleva, M., Cardone, M. H., and Andreeff, M. (2013). BH3 profiling discriminates response to cytarabine-based treatment of acute myelogenous leukemia. Molecular cancer therapeutics 12, 2940–2949.

Pon, J. R., Wong, J., Saberi, S., Alder, O., Moksa, M., Grace Cheng, S. W., Morin, G. B., Hoodless, P. A., Hirst, M., and Marra, M. A. (2015). MEF2B mutations in non-Hodgkin lymphoma dysregulate cell migration by decreasing MEF2B target gene activation. Nature communications 6, 7953.

Schuback, H. L., Arceci, R. J., and Meshinchi, S. (2013). Somatic characterization of pediatric acute myeloid leukemia using next-generation sequencing. Seminars in hematology 50, 325–332.

Schuler, A., Schwieger, M., Engelmann, A., Weber, K., Horn, S., Muller, U., Arnold, M. A., Olson, E. N., and Stocking, C. (2008). The MADS transcription factor Mef2c is a pivotal modulator of myeloid cell fate. Blood 111, 4532–4541.

Schwieger, M., Schuler, A., Forster, M., Engelmann, A., Arnold, M. A., Delwel, R., Valk, P. J., Lohler, J., Slany, R. K., Olson, E. N., and Stocking, C. (2009). Homing and invasiveness of MLL/ENL leukemic cells is regulated by MEF2C. Blood 114, 2476–2488.

Shulman, J. M., Benton, R., and St Johnston, D. (2000). The Drosophila homolog of C. elegans PAR-1 organizes the oocyte cytoskeleton and directs oskar mRNA localization to the posterior pole. Cell 101, 377–388.

Somervaille, T. C., and Cleary, M. L. (2006). Identification and characterization of leukemia stem cells in murine MLL-AF9 acute myeloid leukemia. Cancer cell 10, 257–268.

Stehling-Sun, S., Dade, J., Nutt, S. L., DeKoter, R. P., and Camargo, F. D. (2009). Regulation of lymphoid versus myeloid fate 'choice' by the transcription factor Mef2c. Nature immunology 10, 289–296.

Timm, T., Balusamy, K., Li, X., Biernat, J., Mandelkow, E., and Mandelkow, E. M. (2008). Glycogen synthase kinase (GSK) 3beta directly phosphorylates Serine 212 in the regulatory loop and inhibits microtubule affinity-regulating kinase (MARK) 2. The Journal of biological chemistry 283, 18873–18882.

Wang, W., Org, T., Montel-Hagen, A., Pioli, P. D., Duan, D., Israely, E., Malkin, D., Su, T., Flach, J., Kurdistani, S. K., et al. (2016). MEF2C protects bone marrow B-lymphoid progenitors during stress haematopoiesis. Nature communications 7, 12376.

Ying, C. Y., Dominguez-Sola, D., Fabi, M., Lorenz, I. C., Hussein, S., Bansal, M., Califano, A., Pasqualucci, L., Basso, K., and Dalla-Favera, R. (2013). MEF2B mutations lead to deregulated expression of the oncogene BCL6 in diffuse large B cell lymphoma. Nature immunology 14, 1084–1092.

Yu, Y. T., Breitbart, R. E., Smoot, L. B., Lee, Y., Mahdavi, V., and Nadal-Ginard, B. (1992). Human myocyte-specific enhancer factor 2 comprises a group of tissue-restricted MADS box transcription factors. Genes & development 6, 1783–1798.

Zahreddine, H. A., Culjkovic-Kraljacic, B., Assouline, S., Gendron, P., Romeo, A. A., Morris, S. J., Cormack, G., Jaquith, J. B., Cerchietti, L., Cocolakis, E., et al. (2014). The sonic hedgehog factor GLI1 imparts drug resistance through inducible glucuronidation. Nature 511, 90–93.

Zheng, R., Klang, K., Gorin, N. C., and Small, D. (2004). Lack of KIT or FMS internal tandem duplications but co-expression with ligands in AML. Leukemia research 28, 121–126.

Zhou, H., Di Palma, S., Preisinger, C., Peng, M., Polat, A. N., Heck, A. J., and Mohammed, S., 2013 Toward a comprehensive characterization of a human cancer cell phosphoproteome. Journal of proteome research 12, 260–271.

Zhou, H. S., Carter, B. Z., and Andreeff, M. (2016). Bone marrow niche-mediated survival of leukemia stem cells in acute myeloid leukemia: Yin and Yang. Cancer biology & medicine 13, 248–259.

Zhu, B., and Gulick, T. (2004). Phosphorylation and alternative pre-mRNA splicing converge to regulate myocyte enhancer factor 2C activity. Molecular and cellular biology 24, 8264–8275.

Zuber, J., Radtke, I., Pardee, T. S., Zhao, Z., Rappaport, A. R., Luo, W., McCurrach, M. E., Yang, M. M., Dolan, M. E., Kogan, S. C., et al. (2009). Mouse modelsofhuman AML accuratelypredict chemotherapyresponse. Genes & development 23, 877–889.

Zuber, J., Rappaport, A. R., Luo, W., Wang, E., Chen, C., Vaseva, A. V., Shi, J., Weissmueller, S., Fellmann, C., Taylor, M. J., et al. (2011). An integrated approach to dissecting oncogene addiction implicates a Myb-coordinated self-renewal program as essential for leukemia maintenance. Genes & development 25, 1628–1640.

## SUPPLEMENTAL REFERENCES

Cancer Genome Atlas Research, N. (2013). Genomic and epigenomic landscapes of adult de novo acute myeloid leukemia. N Engl J Med, 368, 2059–2074.

Hu, Y. & Smyth, G. K. (2009). ELDA: extreme limiting dilution analysis for comparing depleted and enriched populations in stem cell and other assays. J Immunol Methods, 347, 70–78.

Kitagawa, D., Gouda, M., Kirii, Y., Sugiyama, N., Ishihama, Y., Fujii, I., Narumi, Y., Akita, K. & Yokota, K. (2012). Characterization of kinase inhibitors using different phosphorylation states of colony stimulating factor-1 receptor tyrosine kinase. J Biochem, 151, 47–55.

Krivtsov, A. V., Twomey, D., Feng, Z., Stubbs, M. C., Wang, Y., Faber, J., Levine, J. E., Wang, J., Hahn, W. C., Gilliland, D. G., et al. (2006). Transformation from committed progenitor to leukaemia stem cell initiated by MLL-AF9. Nature, 442, 818–822.

Meerbrey, K. L., Hu, G., Kessler, J. D., Roarty, K., Li, M. Z., Fang, J. E., Herschkowitz, J. I., Burrows, A. E., Ciccia, A., Sun, T., et al. (2011). The pINDUCER lentiviral toolkit for inducible RNA interference in vitro and in vivo. Proc Natl Acad Sci U S A, 108, 3665–3670.

Naviaux, R. K., Costanzi, E., Haas, M. & Verma, I. M. (1996). The pCL vector system: rapid production of helper-free, high-titer, recombinant retroviruses. J Virol, 70, 5701–5705.

Wang, H., Yang, H., Shivalila, C. S., Dawlaty, M. M., Cheng, A. W., Zhang, F. & Jaenisch, R. (2013). One-step generation of mice carrying mutations in multiple genes by CRISPR/Cas-mediated genome engineering. Cell, 153, 910–918.

Wefers, B., Ortiz, O., Wurst, W. & Kuhn, R. (2014). Generation of targeted mouse mutants by embryo microinjection of TALENs. Methods, 69, 94–101.

Yang, X., Boehm, J. S., Yang, X., Salehi-Ashtiani, K., Hao, T., Shen, Y., Lubonja, R., Thomas, S. R., Alkan, O., Bhimdi, T., et al. (2011). A public genome-scale lentiviral expression library of human ORFs. Nat Methods, 8, 659–661.

Zhu, N., Chen, M., Eng, R., Dejong, J., Sinha, A. U., Rahnamay, N. F., Koche, R., Al-Shahrour, F., Minehart, J. C., Chen, C. W., et al. (2016). MLL-AF9-and HOXA9-mediated acute myeloid leukemia stem cell self-renewal requires JMJD1C. J Clin Invest, 126, 997–1011.

